# Human labour pain is influenced by the voltage-gated potassium channel K_V_6.4 subunit

**DOI:** 10.1101/489310

**Authors:** Michael C. Lee, Michael S. Nahorski, James R.F. Hockley, Van B. Lu, Gillian Ison, Luke A. Pattison, Gerard Callejo, Kaitlin Stouffer, Emily Fletcher, Christopher Brown, Ichrak Drissi, Daniel Wheeler, Patrik Ernfors, David Menon, Frank Reimann, Ewan St John Smith, C. Geoffrey Woods

## Abstract

We sought genetic effects on labour pain by studying healthy women who did not request analgesia during their first delivery. Extensive sensory and psychometric testing were normal in these women, except for significantly higher cuff-pressure pain. We found an excess of heterozygotes carrying the rare allele of SNP rs140124801 in *KCNG4*. The rare variant K_V_6.4-Met419 exerts a dominant negative effect and cannot modulate the voltage-dependence of K_V_2.1 inactivation because it fails to traffic to the plasma membrane. *In vivo*, we observed *Kcng4* (K_V_6.4) expression in 40% of retrograde labelled mouse uterine sensory neurones, all of which expressed K_V_2.1, and over 90% expressed nociceptor genes *Trpv1* and *Scn10a*. In neurones overexpressing K_V_6.4-Met419, the voltage-dependence of inactivation for K_V_2.1 is more depolarised compared to neurones overexpressing K_V_6.4. Finally, K_V_6.4-Met419 overexpressing neurones have a higher action potential threshold. We conclude K_V_6.4 can influence human labour pain by modulating the excitability of uterine nociceptors.

## Introduction

All eutherians (placental mammals) experience contraction of the uterus and discomfort during parturition. Whilst this discomfort is universal in eutherians, it appears to be most marked in humans (*1*). The severity of labour pain is considered a consequence of positive sexual selection in modern humans (with females seeking the cleverest mate), which has led to the human brain (and head) being three times the relative size of our nearest primate relatives (*2*). Despite neoteny (birth of offspring in a relatively immature state), this imposes a need to deliver a large neonatal head through the birth canal causing labour pain (*3*). While labour pain is clearly linked to uterine contractions and cervical distension, the generation of this visceral signal and the sensory afferents involved are poorly understood (*4*).

Although there are well-established ethnic, social and cultural factors that influence the experience and expression of pain during labour (*5*), broader genetic effects on labour pain may also exist. For example, women with the very rare Mendelian disorder Congenital Insensitivity to Pain due to bi-allelic non-functional mutations in *SCN9A* (OMIM: 243000) do not report labour pain or require analgesics during labour (*6*). *SCN9A* encodes for the voltage-gated sodium channel Na_V_1.7, expressed selectively in nociceptive and autonomic neurones, and mutations in *SCN9A* have well-documented roles in causing extremely painful or painless phenotypes (*7*). The painlessness conferred by loss of function *SCN9A* mutations is clearly maladaptive and can be associated with severe injury during human parturition (*8*).

Our aim here was not to discover very rare Mendelian mutations that cause extreme painlessness, for example, congenital insensitivity to pain. Instead, the genetic analyses employed here are optimized for investigation of phenotypes that require both an environmental trigger and genetic predisposition, that will not appear to have a Mendelian inheritance pattern, unless the triggering event is frequent (*9*). This approach is suited to the study of labour pain, which may be considered nociceptive in nature, with parturition serving as visceral stimulus. We sought to identify functional SNP alleles that are over- or under-represented in a cohort of women who did not request or use analgesics that were available and offered to them during labour: an observable behavioural phenotype that is considered highly unusual in hospital maternity units in the United Kingdom, particularly for the spontaneous delivery of term nulliparous women. Quantitative sensory testing, performed with our study cohort, suggest a general increase in pain thresholds and tolerance when compared to controls, but only the increase in cuff-pressure pain threshold survived statistical significance after adjustment for multiple comparisons. We next assessed the allele frequencies of all (genome-wide) protein changing single nucleotide polymorphisms (SNPs) in these women compared to population frequencies. We found that the voltage-gated potassium channel (K_V_) modifier *KCNG4* (K_V_6.4) SNP rs140124801 rare allele c.1255G>A p.(Val419Met) was over-represented. Finally, we demonstrate effects of this rare K_V_6.4-Met419 variant on sensory neurone excitability, and hence reveal a mechanism through which uterine nociception, and hence labour pain, can be attenuated in humans.

## Results

### Identifying women who did not require analgesics during labour as nulliparous parturients: the test cohort

1029 potential cases were identified from 8 maternity units in the United Kingdom over a three-year period. Each potential case was invited to contact researchers, as chronologically ascertained. 383 women responded and were screened via telephone (Figure S1A). Key inclusion criteria were: healthy Caucasian women who experienced term (beyond 37-week gestation) and spontaneous vaginal delivery as nulliparous parturients without any use or request for any form of systemic or regional analgesia (spinal or epidural). We excluded women who had major disease or co-morbidities that are known to influence labour pain or pain in general. 189 women met the full eligibility criteria (Table S1), returned written consent, and donated either 10 ml of blood (collected at their local hospital) or 2 ml of saliva sent via post, from which DNA was extracted.

Of the women who donated DNA, 39 consented for a subsequent study of psychometrics and quantitative sensory testing. These women comprised a subset of the genetic discovery cohort for a case-controlled study (Figure S1B). For the control cohort, we recruited 33 women who were matched in age at delivery of the firstborn and location of maternity service, but who used analgesics during labour and delivery of their firstborn (Table S1). There were no significant differences in the means of new-born weight or head circumference between test and control cohorts (Table 1).

**Table 1.** Key characteristics of test cohort comprising women who did not request or require analgesics during nulliparous term spontaneous labour and controls who did. n, number of participants; SD, standard deviation; * Sidak’s correction; CI5-CI95, 5-95% confidence interval; # missing clinical record; ## equipment failure or unavailable; SFMPQ, short-form McGill’s Pain Questionnaire.

| Characteristics<br>(at delivery of first-born) | Test cohort |  |  | Control cohort |  |  | P | P | CI5 | CI95 |
| --- | --- | --- | --- | --- | --- | --- | --- | --- | --- | --- |
|  | n | mean | SD | n | mean | SD | unadjusted | adjusted * |  |  |
| Age (years) | 39 | 32.83 | 4.18 | 33 | 31.94 | 3.98 | 0.33 |  | -2.73 | 0.93 |
| Head circumference of newborn (cm) | #26 | 34.00 | 0.98 | +24 | 34.46 | 0.97 | 0.10 |  | -0.10 | 1.01 |
| Weight of newborn (g) | 38 | 3362 | 434.1 | 33 | 3384 | 419.2 | 0.83 |  | -180.90 | 224.76 |
| Characteristics<br>(at research visit) | Test cohort |  |  | Control cohort |  |  | P | P | CI5 | CI95 |
|  | n | mean | SD | n | mean | SD | unadjusted | adjusted * |  |  |
| Age (years) | 39 | 36.26 | 4.18 | 33 | 36.45 | 4.11 | 0.62 |  | -1.48 | 2.46 |
| Upper arm diameter at assessment (cm) | 39 | 28.54 | 3.60 | 33 | 29.23 | 3.63 | 0.43 |  | -1.03 | 2.41 |
| **Sensory and pain thresholds** |  |  |  |  |  |  |  |  |  |  |
| Cold detection (°C) | 39 | 30.45 | 0.93 | 33 | 30.35 | 0.95 | 0.79 |  | -0.42 | 0.25 |
| Warm detection (°C) | 39 | 34.43 | 0.99 | 33 | 34.97 | 0.87 | 0.002 | 0.012 | 0.28 | 1.05 |
| Cuff pressure detection (mmHg) | 39 | 28.44 | 7.79 | 33 | 27.10 | 8.38 | 0.51 |  | -5.00 | 1.33 |
| Cold pain (°C) | 39 | 11.64 | 8.26 | 33 | 16.88 | 9.03 | 0.02 | 0.114 | 1.17 | 9.73 |
| Heat pain (°C) | 39 | 44.08 | 2.85 | 33 | 42.36 | 3.40 | 0.018 | 0.103 | -2.92 | -0.27 |
| Cuff pressure pain (mmHg) | 39 | 166.7 | 54.74 | 33 | 113.03 | 42.96 | 0.00002 | 0.00012 | -77.03 | -30.13 |
| **Pain tolerance (cold immersion)** |  |  |  |  |  |  |  |  |  |  |
| Pre-immersion hand temperature (°C) | 34 | 30.46 | 1.95 | 33 | 30.82 | 1.66 | 0.42 |  | -0.53 | 1.24 |
| Post-immersion hand temperature (°C) | ##33 | 17.92 | 4.72 | 33 | 20.51 | 3.54 | 0.02 | 0.12 | 0.40 | 4.60 |
| Latency to hand withdrawal (s) | ##36 | 77.03 | 71.82 | 33 | 44.11 | 55.73 | 0.03 | 0.14 | -38.0 | -0.0000 |
| Peak pain occurrence (0-100mm) | ##35 | 80.19 | 27.39 | 33 | 79.04 | 28.99 | 0.71 |  | -5.50 | 4.00 |
| Peak pain intensity (0-100mm) | ##35 | 54.29 | 17.26 | 33 | 65.82 | 13.20 | 0.004 | 0.02 | 3.20 | 18.1 |
| SFMPQ (sensory) | 36 | 8.47 | 3.82 | 33 | 10.97 | 4.00 | 0.010 | 0.049 | 0.62 | 4.38 |
| SFMPQ (affective) | 36 | 1.00 | 1.53 | 33 | 1.24 | 1.35 | 0.26 |  | -0.00002 | 0.99995 |

### Cognitive and emotional function is normal in the test cohort

Psychometrics, comprising validated questionnaires and computerized cognitive assessments, were employed to quantify mood, beliefs and personality traits that can influence pain in experimental or clinical settings. The questionnaires included were: Hospital Anxiety and Depression Scale (HADS) (*10*), Pain Catastrophizing Scale (PCS) (*11*), Multidimensional Health Locus of Control Scale (MHLC) (*12*) and Life Orientation Test-Revised (LOTR) (*13*). Computerized cognitive assessments were implemented in CANTAB® (Cambridge Cognition, UK) (*14*). There were no significant differences in psychological or cognitive measures between control and test cohorts (Table S2).

### Experimental pain thresholds and tolerance are increased in the test cohort

Next, we quantified sensory detection and pain thresholds to cold, heat and mechanical pressure. Thermal stimuli were delivered using a skin thermode applied to the forearm. Mechanical pressure was exerted via compression of upper arm by a sphygmomanometer cuff. There were no significant differences in the detection thresholds of cold or cuff-pressure in the test and control cohorts to suggest sensory deficits or impairments pertaining to those stimuli in the test cohort (Table 1, Figure S2A). Warmth detection thresholds were very slightly but significantly lower in the test cohort compared to controls (0.54 °C difference) but all individual values fell within established norms for the general population (*15*).

The test cohort had increased pain thresholds to heat, cold and cuff-pressure compared to controls at an unadjusted significance level of *P*<0.05 (Figure S2A). There was a very striking increase of over 50 mmHg in the cuff-pressure pain threshold (*P* = 0.00002, uncorrected; *P* = 0.00012, Sidak’s correction) (Table 1), suggesting that this characteristic might be relevant to the lack of analgesic requirement during nulliparous labour in the test cohort.

During testing for tolerance to pain from the immersion of hand in cold water (3 °C), when compared to controls, the test cohort showed increased hand withdrawal latency (*P* = 0.03, uncorrected), lower post-immersion skin temperatures (*P* = 0.02, uncorrected), and lower peak intensity of pain on the 100 mm Visual Analogue Scale (VAS) (*P* = 0.004, uncorrected; *P* = 0.02, Sidak’s correction) on later assessment (Figure S2B). The Short-Form McGill Pain questionnaire (*16*) revealed lower scores (*P* = 0.01, uncorrected; *P* = 0.049, corrected) for the sensory descriptors for the test group. There was no between-group difference in scores related to the affective aspects of the experimentally induced pain experienced (*P* = 0.26). These individual results do not survive statistical correction for multiple comparisons but suggest that overall the test cohort of women who did not request analgesia during nulliparous parturition showed greater tolerance to pain when compared to controls.

### The rare allele of rs140124801 in *KCNG4* is over-represented in the test cohort

In 158 of the 189 women who did not require analgesics during their first labour, we obtained enough high-quality DNA for molecular genetic analysis (Figure S1). The chronologically first 100 such women (by date of banking DNA) constituted a discovery cohort (Figure 1A); the next 58 women constituted our replication cohort. Those in the discovery cohort each had exome sequencing, from which we used the bam and bam.bai files for genome wide SNP allele frequency assessment using the fSNPd programme (*9*). The replication cohort of 58 were assessed only for SNP rs140124801 alleles using Sanger sequencing of genomic DNA.

**Figure 1.**
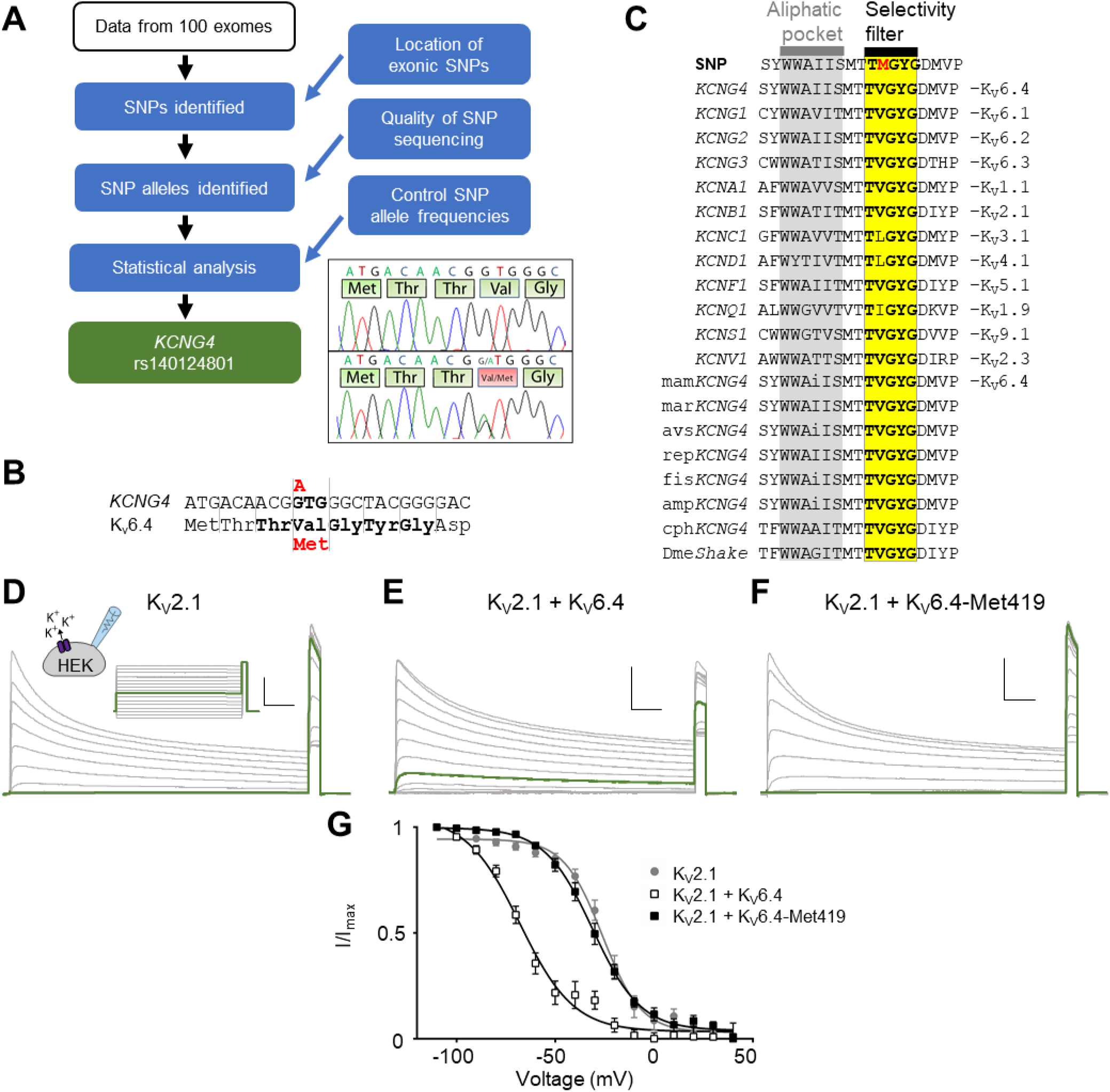
Molecular genetics of *KCNG4* SNP rs140124801, and analysis of K_V_2.1 inactivation properties (A) Summary of the genetic analysis. The resultant finding is of the SNP rs140124801 in *KCNG4*. Inset: electrophoretograms showing the alleles. (B) The nucleotide sequence of the SNP rs140124801 (NM_1.NM_172347.2) showing the altered GTG codon (in bold), and the rare allele (in red). Amino acids 416 to 423 of K_V_6.4 (NP_758857.1) are shown below their nucleotide codons. The selectivity filter is in bold, and the wild type Val-419 shown above Met-419. (C) Evolutionary conservation of human K_V_6.4 positions 408 to 426: rs140124801 alleles, representative proteins of each human K_V_ class, and of K_V_6.4 in vertebrates. Invariant amino acids are capitalized. The selectivity filter TVGYG in yellow, and conserved aliphatic region in grey. Representative current recordings to determine K_V_2.1 (D), K_V_2.1/KV6.4 (E), K_V_2.1/K_V_6.4-Met419 (F) steady-state inactivation properties. The applied voltage protocol is illustrated above (D). Vertical scale bar is 10 nA, horizontal scale bar is 0.5 s. Green traces indicate currents recorded during the −40 mV conditioning step. (G) Voltage-dependence of steady-state inactivation of K_V_2.1 (grey filled circles, n = 9), K_V_2.1/K_V_6.4 (white squares, n = 12), and K_V_2.1/K_V_6.4-Met419 (black squares, n = 15). Solid lines represent the Boltzmann fitted curves.

Our discovery cohort analysis identified one ion channel SNP where the allele frequency was altered compared to reference (Figure 1A). The rare allele of rs140124801 in *KCNG4* was over-represented, being found in 3 instances, whereas 0.7 instances were expected (q=<0.05, FDR corrected). We examined the individual exome results using the Integrated Genome Viewer (https://software.broadinstitute.org/software/igv/) and found that 3 individuals were heterozygous for the rare allele and confirmed this by Sanger sequencing. In the replication cohort, we found one further rare SNP rs140124801 heterozygote. For the total cohort of 158 women not requiring analgesia during their first delivery, there were 4 heterozygotes carrying the rs140124801 rare allele compared to an expected 1.1 (Chi-squared two tail with Yates correction = 4.779, *P* = 0.0288; Figure S1A).

In case-controlled studies, we further explored whether 3 of the individuals who possess the rare *KCNG4* allele had significantly different experimental pain thresholds to those who did not (n=69, Figure S1B). We investigated pain thresholds for heat, cold and cuff-pressure, and found that the rare *KCNG4* allele was associated with a significantly increased cuff-pressure pain threshold (*P* = 0.0029, uncorrected; *P* = 0.009, Sidak’s correction, Table S3). Although the sample size here is very small because of the rarity of the *KCNG4* allele being examined, the finding suggests that an effect of this rare-allele is to increase experimental cuff-pressure pain threshold in humans. The experimental cuff-pressure pain remains significantly increased in the test cohort (even with the 3 rare-allele cases excluded, when compared with the control group (*P* = 0.0029, uncorrected; *P* = 0.009, Sidak’s correction, Table S3) suggesting that cuff-pressure pain threshold might be relevant to the labour pain. Whilst, there are clearly other reasons for increased cuff-pressure pain threshold in those cases who do not carry the rare *KCNG4* allele, these data suggest that the rare-allele of *KCNG4* may be related to the lack of analgesic requirement for the 3 cases we identified in this study.

### The rare allele of rs140124801 in *KCNG4* introduces a p.Val419Met change in the highly conserved K^+^ selectivity filter consensus sequence

The rare allele of rs140124801 in *KCNG4* causes the mis-sense change p.Val419Met encoding the voltage-gated potassium channel K_V_6.4 (from here on referred to as K_V_6.4-Met419; Figure 1A-B). Voltage-gated potassium channels are tetrameric complexes with each subunit having six transmembrane domains (S1-S6). K_V_6.4 is a member of the electrically silent group of K_V_ subunits, which cannot form functional plasma membrane-expressed homotetramers, but instead act as modulators of K_V_2 subunits (*17*). Indeed, K_V_6.4 is known to heterotetramerise with K_V_2.1 in a 1:3 stoichiometry (*18*). Valine 419 is in the pore forming S5-S6 linker and is part of the highly conserved K^+^ selectivity filter consensus sequence (T**V**GYG) (Figure 1C), in which the equivalent position is always occupied by a branched chain amino acid. Whilst originally thought to be relatively rigid, this structure is also involved in open-pore or C-type inactivation, as subtle rearrangements block the conductive path of K^+^ ions (*19*). The molecular mechanisms underlying K_V_2.1 channel inactivation remain unclear, but include interference with the predominant U-type inactivation from closed states (*20*) by the K_V_2.1-E352Q mutation (*21*) or, in the case of K_V_6.4/K_V_2.1 heterotetrameric channels, treatment with 4-aminopyridine (*22*), unveiled C-type inactivation. K_V_6.4/K_V_2.1 heterotetramers display a hyperpolarizing shift in the voltage-dependence of inactivation compared to the K_V_2.1 homotetramers, which has been linked to differences in the voltage sensing S4 segment (*23*), but additional contributions from the S5-S6 linker have not been excluded. It seemed likely that rs140124801 might affect K^+^-selectivity and/or inactivation. We therefore studied the electrophysiological properties of K_V_6.4-Met419 in complex with K_V_2.1 compared to the most frequent *KCNG4* allele that possesses a valine at position 419 (K_V_6.4) in complex with K_V_2.1.

### The p.Val419Met change in KV6.4 impairs function of KV2.1 heterotetramers

We used HEK293 cells as a heterologous expression system that does not express significant endogenous K_V_ currents (Figure S3). As expected, over-expression of K_V_6.4 or K_V_6.4-Met419 alone did not produce measurable K^+^ currents (Figure S3E). However, in cells expressing K_V_2.1 alone, outward currents were observed that were activated by potentials more positive than −40 mV and displayed a slow inactivation (Figure S3A). Co-expression of K_V_2.1 with K_V_6.4 produced outward currents with similar kinetics (Figure S3D), but we observed a small shift in the voltage of half-maximal activation (V_0.5_ act) to more negative potentials. This shift was not observed when K_V_6.4-Met419 was co-expressed with K_V_2.1 (Figure S3D). The current amplitude generated was similar between wild-type K_V_6.4 or K_V_6.4-Met419 co-expressed with K_V_2.1 (Figure S3E) showing that expression of K_V_6.4-Met419 does not negatively regulate maximal current flux, over wild-type K_V_6.4, a factor that would impact sensory neurone excitability (Figure S3E). The slope factors of the Boltzmann fits did not significantly differ between the 3 groups (K_V_2.1: k = 9.5 ± 0.8, n = 13; K_V_2.1 + K_V_6.4: k = 15.9 ± 1.7, n = 14; K_V_2.1 + K_V_6.4-Met419: k = 11.0 ± 0.8, n = 13; one-way ANOVA, P > 0.05). Furthermore, the reversal potential was not significantly different between the groups (Figure S3F).

Similar to previous reports (*23*), co-expression of K_V_6.4 resulted in a large hyperpolarising shift in the voltage-dependence of inactivation by ∼30 mV compared to K_V_2.1 homomeric currents (Figure 1D, E & G). This hyperpolarising shift was not observed when K_V_2.1 was co-expressed with K_V_6.4-Met419 (Figure 1F & G). There was however no significant difference in the slope factor of inactivation curves between the three groups (K_V_2.1: k = 9.8 ± 1.4, n = 9; K_V_2.1 + K_V_6.4: k = 13.6 ± 2.4, n = 12; K_V_2.1 + K_V_6.4-Met419: k = 12.2 ± 1.2, n = 15; Kruskal-Wallis, *P* > 0.7), or in their time courses of recovery from inactivation (Figure S3G). These data suggest a loss of K_V_6.4 function as a result of the p.Val419Met mutation.

### K_V_6.4-Met419 does not traffic with K_V_2.1 to the plasma membrane

As discussed above, K_V_6.4 forms heterotetramers with K_V_2.1 with altered biophysical properties compared to homotetrameric K_V_2.1 channels (*24*) (Figure 1D-G, Figure S3). In addition, K_V_6.4 is retained in the endoplasmic reticulum in the absence of K_V_2.1, requiring the expression of K_V_2.1 for trafficking to the cell membrane (*25*). We thus tested whether the p.Val419Met alteration might affect the trafficking of K_V_6.4. For this, K_V_6.4 was cloned into a pcDNA3 based vector containing a CMV-polioIRESmCherry expression cassette, tagged with HA and the p.Val419Met alteration introduced. K_V_2.1 had been previously cloned into the pCAGGS-IRES2-nucEGFP which displays nuclear GFP signal upon transfection. To assess membrane localisation, HEK293 cells were co-transfected with both K_V_2.1 and K_V_6.4, stained for HA-tagged K_V_6.4, with co-expressing cells identified by both mCherry and nuclear GFP signal. K_V_6.4 was retained within the cytoplasm in the absence of K_V_2.1 expression but displayed a striking shift to the cell membrane upon co-transfection with K_V_2.1 (Figure 2A). There was no appreciable difference in the localization of K_V_6.4-Met419 in the absence of K_V_2.1, but in the presence of K_V_2.1 and in contrast to the wild-type protein, K_V_6.4-Met419 was retained intracellularly and showed no membrane localization (Figure 2A). Importantly, expression of K_V_6.4-Met419 in HEK293 cells showed only a modest reduction in steady-state stability compared with wild-type K_V_6.4, and this was not affected by co-expression with K_V_2.1 (Figure 2B-C).

**Figure 2.**
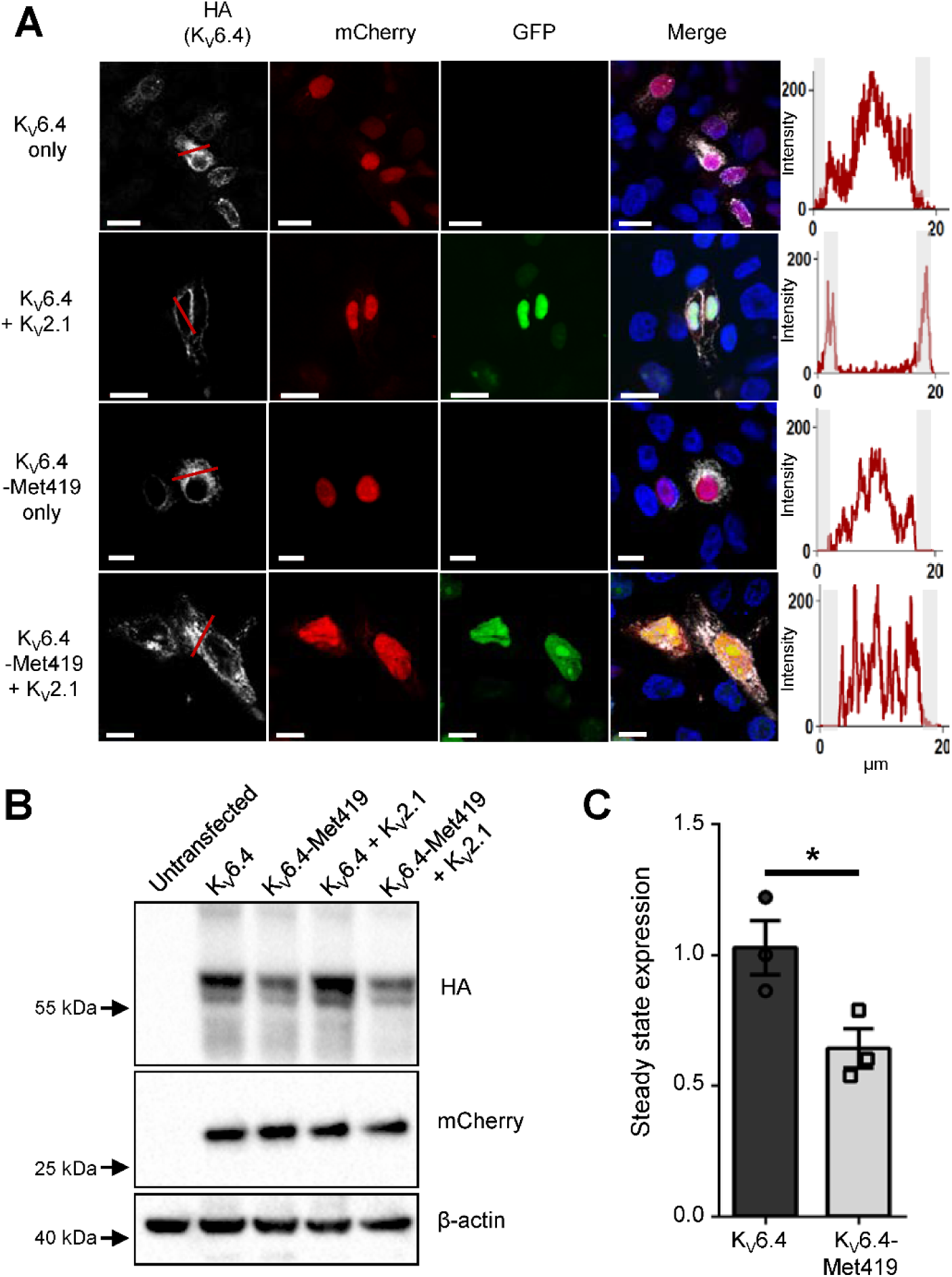
p.Val419Met blocks K_V_6.4 from reaching the plasma membrane independent of changes in steady-state expression (A) Immunofluorescence analysis of K_V_6.4 localization. In the absence of K_V_2.1, K_V_6.4 was retained in the cytoplasm (white channel, top panel), and was trafficked to the cell membrane in the presence of K_V_2.1 (white channel, 2nd panel down, white arrow). In contrast, HA-tagged K_V_6.4-Met419 did not localize to the cell membrane in either the absence or presence of K_V_2.1 expression (white channels in the 3rd and 4th panel down, white arrow). Expression of K_V_2.1 is demonstrated by presence or absence of green nuclei, expression of K_V_6.4 is displayed directly by HA tag in the white channel and expression of the IRES vector expressing K_V_6.4 is displayed by presence of mCherry signal in the red channel. Graphs adjacent to each panel display the intensity of K_V_6.4 HA signal along the red line in each respective white channel; note membrane localized peaks only in K_V_6.4 when co-expressed with K_V_2.1. Scale bars indicate 10 µm (B) HA-tagged K_V_6.4 was transiently expressed in the presence or absence of K_V_2.1. There was a modest reduction in steady state stability for K_V_6.4-Met419 compared with K_V_6.4. (C) Stability assessed by densitometry of HA compared with mCherry as a control of transfection efficiency. Unpaired t-test (P = 0.04).

### K_V_6.4 is expressed in nociceptors that innervate the uterus

Altered K_V_ function produces dramatic effects upon sensory neurone excitability; K_V_7 openers (*26*) and K_V_2 inhibitors (*27*) decrease and increase sensory neurone excitability respectively. We hypothesised that expression of K_V_6.4-Met419 within sensory neurones innervating the uterus would alter neuronal excitability and contribute to the impaired nociception. We first investigated the expression of *Kcng4* and *Kcnb1* in mouse uterine sensory neurones using single-cell qRT-PCR of sensory neurones retrogradely labelled with fast blue from the uterus (Figure 3A). Sensory innervation of the mouse uterus possesses two distinct peak densities within thoracolumbar (TL) and lumbosacral (LS) spinal segments (*28*). As such, fast blue-positive uterine sensory neurones were collected from dorsal root ganglia (DRG) isolated from vertebrae levels T12-L2 and L5-S2. These had an average cell diameter of 31.0 ± 0.7 µm (n = 89), which is in broad agreement with studies investigating sensory neurones innervating the uterus and other visceral organs including the distal colon (*28, 29*). Most uterine neurones expressed *Kcnb1* (TL: 82% [36/44] and LS: 66% [30/45]) and *Kcng4* mRNA was detected in a subset of uterine neurones from both spinal pathways (TL: 43% [19/44] and LS: 24% [11/45]; Figure 3B). The average cycle threshold (CT) value for *Kcng4* expressing neurones was higher than that of *Kcnb1* (27.2 vs. 16.3, Figure S4), which may indicate relative lower expression levels. Importantly, all but one LS neuron co-expressed *Kcng4* with *Kcnb1*, suggesting that these two K_V_ subunits are predominantly present in the same uterine sensory neurone subset. We also assessed the mRNA expression of the nociceptor markers transient receptor potential vanilloid 1 (*Trpv1*) and voltage-gated sodium channel 1.8 (S*cn10a*). In *Kcng4*-positive uterine sensory neurones *Trpv1* mRNA was present in 100 % of TL and 91 % of LS neurones, and *Scn10a* in 95 % of TL and 91 % of LS neurones, suggesting that K_V_6.4 is expressed by a population of neurones capable of transducing noxious stimuli. (Figure 3B).

**Figure 3.**
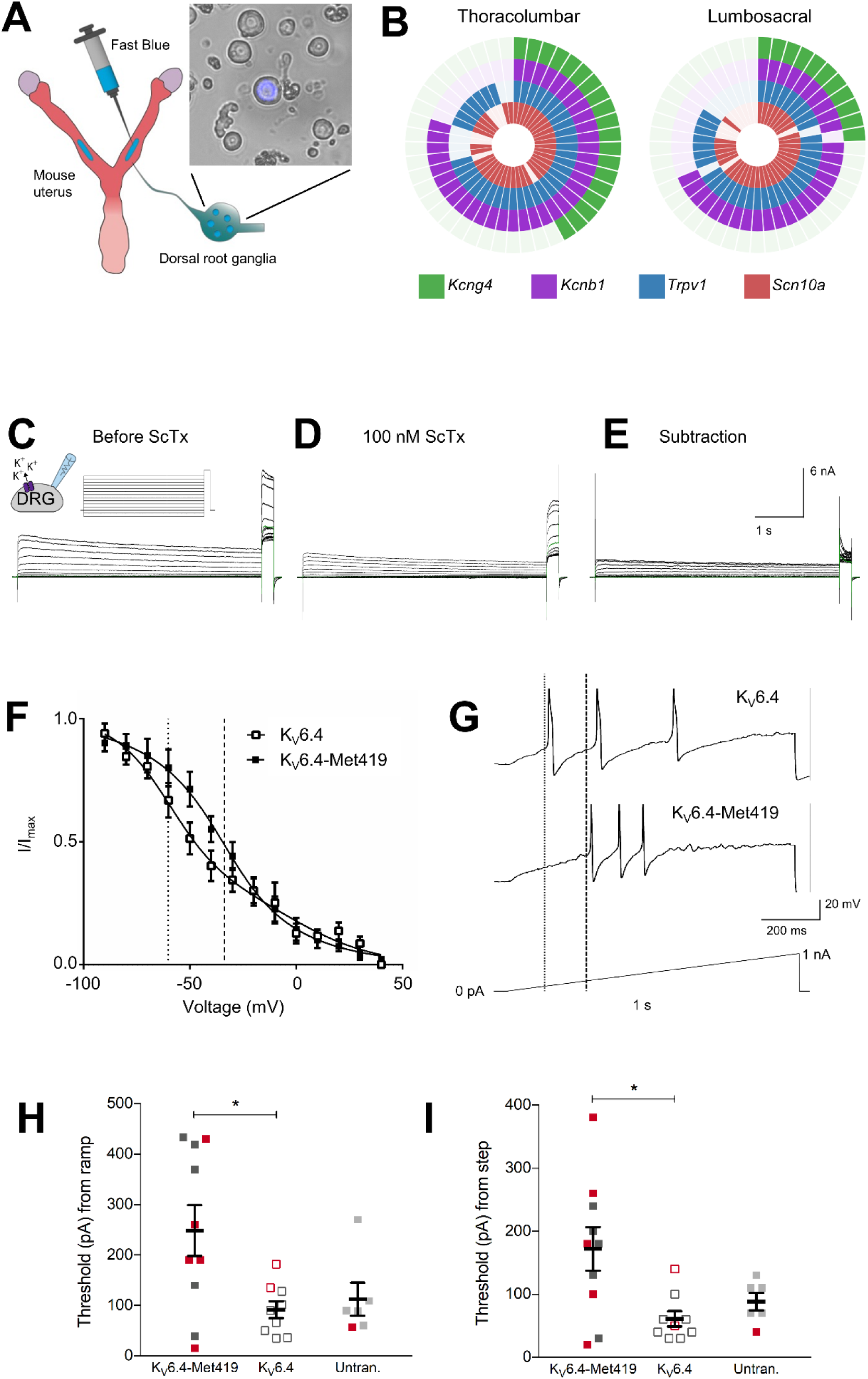
*Kcng4* is coexpressed with *Kcnb1* in mouse uterine sensory neurones and expression of K_V_6.4-Met419 in mouse sensory neurones increases the threshold for action potential discharge, compared to K_V_6.4 (A) Uterine sensory neurones were retrogradely labelled using fast blue and harvested following dissociation. (B) Co-expression analysis of thoracolumbar (T12-L2) and lumbosacral (L5-S2) uterine sensory neurones expressing transcripts for *Kcng4*, *Kcnb1*, *Trpv1* and *Scn10a*. Each segment in the wheel-diagram is representative of a single cell, with a coloured segment signifying positive expression. (C) Representative current recordings to determine the voltage dependence of steady-state inactivation of the stromatoxin-1 (ScTx)-sensitive *I*_K_ elicited by the inset voltage protocol in the absence (C) and presence (D) of 100nM ScTx. Green traces indicate currents recorded during the −40 mV conditioning step. (E) The ScTx-sensitive *I*_K_ was obtained by subtraction of D from C. (F) Inactivation curves for the ScTx-sensitive *I*_K_ for neurones transfected with either K_V_6.4 or K_V_6.4-Met419. Both datasets were fit with a sum of two Boltzmann functions. The midpoints of both the 2nd components of these fits are plotted as either light dashed (K_V_6.4) or heavy dashed (K_V_6.4-Met419) lines. (G) Representative current clamp recordings of neurones of comparable capacitance transfected with either K_V_6.4 or K_V_6.4-Met419 showing action potentials evoked by ramp injection of current (0-1 nA, 1s). The thresholds for action potential discharge are annotated with light dashed (K_V_6.4) or heavy dashed (K_V_6.4-Met419) lines. Summary data of action potential thresholds obtained from neurones transfected with either K_V_6.4 or K_V_6.4-Met419 and untransfected controls obtained via a (H) ramp protocol (0-1 nA, 1s) or (I) step protocol (+10 pA, 50 ms). Red points represent cells that responded to 1 µM capsaicin in voltage clamp mode. Both recordings in Panel G were from cells which were capsaicin responders. Bars indicate mean values, error bars indicate SEM, n = 6-10, * P < 0.05, one-way ANOVA with Bonferroni’s correction for multiple tests.

### K_V_6.4–Met419 causes loss of modulatory function of K_V_2.1 and decreases neuronal excitability in DRG sensory neurones

Given the high co-expression of *Kcng4* with *Kcnb1* in uterine sensory neurones, we next characterized the effect of K_V_6.4 and K_V_6.4–Met419 on sensory neuronal function. We recorded outward delayed rectifier K^+^ currents (*I*_K_) and investigated the effect of transient transfection of either K_V_6.4 or K_V_6.4-Met419 on the stromatoxin-1(ScTx)-sensitive *I*_K_; ScTx is a gating modifier of K_V_2.1, K_V_2.2 and K_V_4.2 which effectively blocks these channels (*30*), as well as K_V_2.1 heterotetramers formed with silent K_V_ subunits (*31*). Through subtraction of *I*_K_ in the presence of ScTx from the total *I*_K_ in the absence of ScTx, we isolated the ScTx-sensitive *I*_K_, which is predominantly dependent on K_V_2 channels (Figure 3C-F). A diverse and heterogenous population of K_V_2 and silent K_V_ subunits is expressed in sensory neurones (*29, 32, 33*) and previous studies suggest that silent K_V_ subunits only heterotetramerise with K_V_2 subunits and not K_V_1, K_V_3 and K_V_4 subunits (*24*). As such, we predicted that wild-type K_V_6.4 heterotetramerisation with K_V_2.1 in sensory neurones would produce functional channels, but with a hyperpolarised shift in the voltage-dependence of inactivation compared to homotetrameric K_V_2.1 channels, as we (Figure 1D-G) and others have observed previously in HEK293 cells (*24*). By contrast, we hypothesised that the K_V_6.4-Met419 subunit would be unable to evoke such a hyperpolarising shift in the voltage-dependence of inactivation.

By transfecting mouse sensory neurones with either K_V_6.4 or K_V_6.4-Met419, we attempted to bias available K_V_2.1 into heterotetramers with K_V_6.4 subunits, thus increasing the probability of recording the contribution of K_V_2.1/K_V_6.4 heterotetramers to ScTx-sensitive *I*_K_. In both K_V_6.4 and K_V_6.4-Met419 experiments, addition of ScTx led to a maximum reduction in the outward K^+^ current at a 20 mV step potential, which did not differ significantly (K_V_6.4, 52.7 ± 3.8 %; K_V_6.4-Met419, 45.1 ± 7.7 %; Student’s t-test, *P* = 0.37; Figure 3C-E). The voltage-dependence of ScTx-sensitive *I*_K_ activation was similar for neurones transfected with K_V_6.4 or K_V_6.4-Met419 subunit (*V*_1/2_ = −5.4 ± 1.8 mV *vs.* −9.8 ± 1.1 mV, and *k* = 8.6 ± 1.5 *vs.* 8.9 ± 0.9, respectively; Figure S5). As observed previously (*33*), the voltage-dependence of ScTx-sensitive *I*_K_ inactivation, for both K_V_6.4 and K_V_6.4-Met419 experiments, was multifactorial and fitted with a sum of two Boltzmann functions. In neurones transfected with K_V_6.4, the midpoint of the first component was −0.8 ± 29.5 mV, which likely correlates with homotetrameric K_V_2.1 currents. The second component possessed a midpoint of inactivation of −60.2 ± 6.6 mV (*n* = 8); a current that is likely a function of heterotetrameric K_V_2/silent K_V_ channels or differentially phosphorylated K_V_2 channels and in line with what others have reported for the second component of *I*_K_ in DRG neurones in the presence of ScTx (*33*). Importantly, expression of K_V_6.4-Met419 led to a significant depolarising shift in the second component of the voltage-dependence of inactivation (−33.8 ± 2.1 mV, *n* = 7, unpaired t-test, *P* = 0.003, Figure 3F), whilst the first component, attributed to homotetrameric K_V_2.1 *I*_K_, remained unchanged (−36.2 ± 3.3 mV, unpaired t-test, *P* = 0.29, Table S4A).

We assessed the functional consequences on neuronal excitability of such a shift in the availability of K_V_2 channels towards more depolarised potentials through current clamp experiments. The threshold for action potential discharge was assessed for neurones transfected with either K_V_6.4 or K_V_6.4-Met419, as well as neurones which exhibited no mCherry fluorescence from cultures exposed to either plasmid (considered untransfected). Neurones transfected with K_V_6.4-Met419 exhibited a higher threshold, than those overexpressing K_V_6.4 or untransfected neurones during injection of a progressively depolarising current, (ramp protocol: 0-1 nA, 1s), however, only the difference between K_V_6.4-Met419 and K_V_6.4 reached statistical significance (K_V_6.4, 91.6 ± 16.7 pA *vs.* K_V_6.4-Met419, 248.6 ± 50.3 pA, ANOVA with Bonferroni multiple comparisons *P* = 0.018; Untran., 112.5 ± 32.5 pA *vs.* K_V_6.4-Met419 248.6 ± 50.3 pA, *P* = 0.087; Figure 3G-H). A higher current was also required to evoke action potentials when threshold was assessed with a step protocol (+10 pA, 50 ms injections, starting at 0 pA). Similarly, only the difference between K_V_6.4 and K_V_6.4-Met419 proved significant (K_V_6.4, 61.1 ± 12.2 pA *vs.* K_V_6.4-Met419, 172.0 ± 34.4 pA, ANOVA with Bonferroni multiple comparisons *P* = 0.012; Untran., 88.3 ± 13.8 pA *vs.* K_V_6.4-Met419 172.0 ± 34.4 pA, *P* = 0.124; Figure 3I). The ability of neurones to respond to capsaicin was also examined to identify putative nociceptors (i.e. those expressing *Trpv1*), but no obvious pattern regarding the subpopulations of nociceptive and non-nociceptive neurones within each group could be observed. Analyses of other action potential parameters revealed no further differences between neurones transfected with either K_V_6.4 construct or untransfected cells (Table S4B). Taken together, these findings demonstrate that sensory neurones expressing K_V_6.4-Met419 are less excitable than those transfected with the K_V_6.4. We thus postulate that uterine primary afferent input into the pain pathway is likely to be reduced in women carrying the rare *KCNG4* SNP rs140124801 allele.

### Heterozygous K_V_6.4-Met419 acts as a dominant negative mutation to abolish wild-type function

The SNP rs140124801 minor allele identified in those healthy women not requiring analgesia during their first labour was always in a heterozygote state. We asked if this heterozygous state has as much of an effect on K_V_2.1 as the homozygous state used in our sub-cellular localisation and electrophysiology studies, or if the effect size was in-between homozygous K_V_6.4 and homozygous K_V_6.4-Met419. Indeed, our findings of reduced labour pain are compatible with the minor allele of rs140124801 having a dominant-negative effect, or a reduced dosage effect, but incompatible if acting as a recessive. K_V_2.1 was co-transfected into HEK293 cells with equimolar concentration of K_V_6.4 and K_V_6.4-Met419, stained for HA-K_V_6.4 and the membrane marker Na^+^/K^+^ ATPase. We found significant co-localisation of K_V_6.4 with Na^+^/K^+^ ATPase at the plasma membrane, but no evidence of trafficking to the cell membrane for either homozygote K_V_6.4-Met419, nor when K_V_6.4 and K_V_6.4-Met419 were co-transfected (Figure 4A-B).

**Figure 4.**
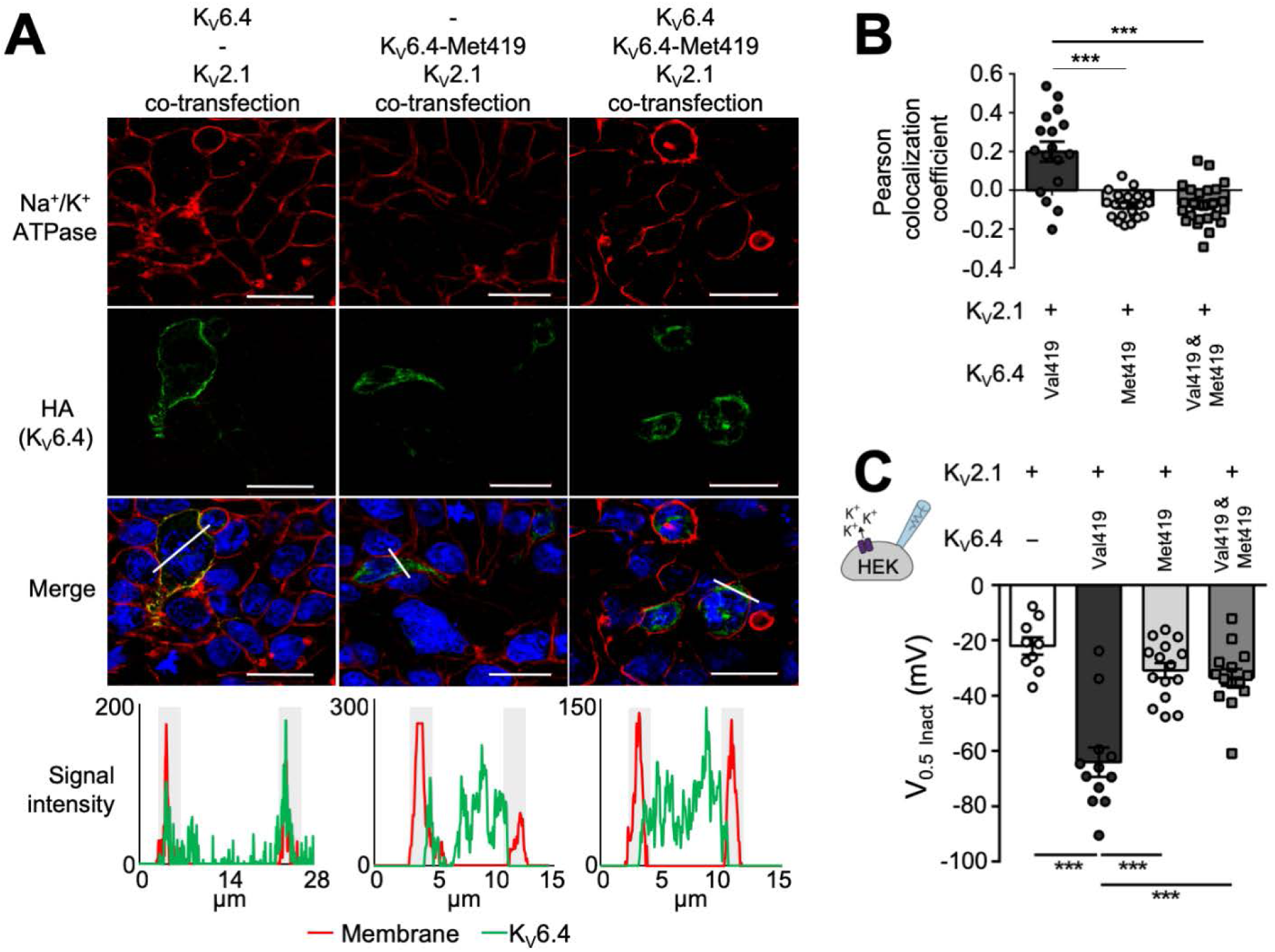
Sub-cellular localization and electrophysiology analysis of the dominant-negative effect of human K_V_6.4-Met419 (A) HEK293 and HeLa cells (separate experiments) were transfected with K_V_2.1 and either wild-type K_V_6.4, K_V_6.4-Met419 or equimolar concentrations of K_V_6.4/K_V_6.4-Met419. Cell membranes were stained with Na^+^/K^+^ ATPase (red channel) and HA-tagged K_V_6.4 (green channel). HA-tagged K_V_6.4 localized to the cell membrane, showing significant co-localization with Na^+^/K^+^ ATPase. Both K_V_6.4-Met419 and K_V_6.4/K_V_6.4-Met419 co-expression showed cytoplasmic retention of K_V_6.4 and no evidence of co-localization with Na^+^/K^+^ ATPase. Graphs below each pane display the profile of signal for membrane and K_V_6.4.HA along the plane of the white line in the merged image. Note red and green signal co-localise in the K_V_6.4 experiment and are distinct in the K_V_6.4-Met419 and heterozygote experiment. (B) Quantification of Pearson’s co-localization co-efficient between K_V_6.4.HA and Na^+^/K^+^ ATPase in each experimental condition. For each condition at least 17 cells were counted from three independent experiments. (C) Voltage of half-maximal inactivation from inactivation protocols shown in Figure 1D-G. Co-expression of both K_V_6.4 and K_V_6.4-Met419 with K_V_2.1 failed to evoke a shift in the voltage-dependence of inactivation. Bars indicate mean values, error bars indicate SEM, n = 9-15, *** *P* < 0.001. Statistics in B and C represent one-way ANOVA with Bonferroni’s multiple comparisons test.

Similarly, co-transfection of equimolar K_V_6.4 and K_V_6.4-Met419 with K_V_2.1 produces electrophysiological properties comparable to transfection of K_V_2.1 only, i.e. the co-expression of the minor allele variant prevented the hyperpolarising shift of the voltage-dependence of inactivation produced by the major allele variant (Figure 4C).

In addition, we investigated whether K_V_6.4-Met419 might affect heterotetramerisation with K_V_2.1. Co-immunoprecipitation experiments in transfected HEK293 cells demonstrate that, unlike K_V_6.4, K_V_6.4-Met419 is unable to bind to K_V_2.1 (Figure S6). When K_V_6.4 is tagged but co-expressed with K_V_6.4-Met419 (untagged), there is notably reduced binding of K_V_6.4 to K_V_2.1 (Figure 5A). Similarly, by immunofluorescence analysis, the presence of untagged K_V_6.4-Met419 suffices to disrupt K_V_6.4 trafficking to the plasma membrane (Figure 5B and 5C).

**Figure 5.**
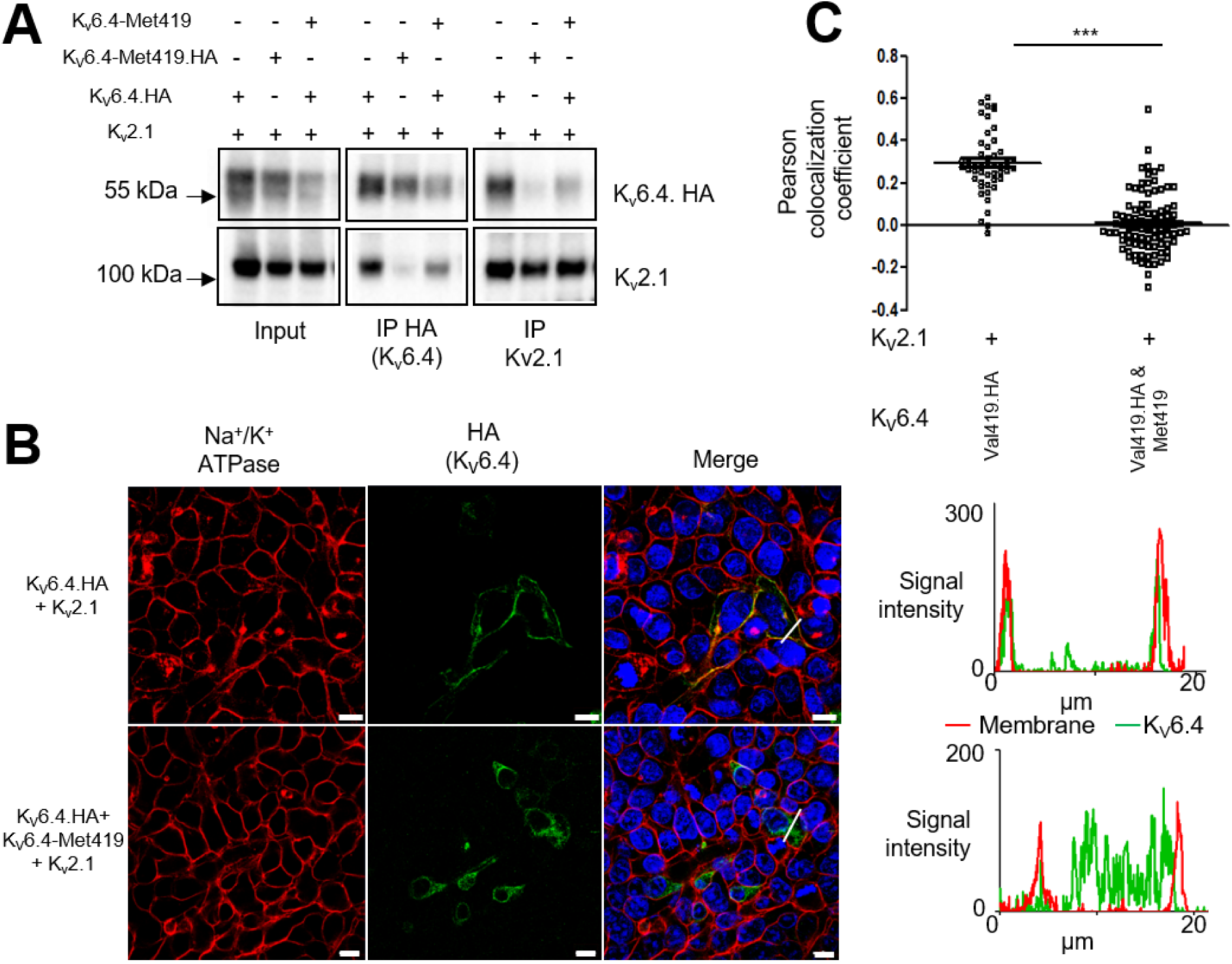
Effects of K_V_6.4-Met419 on K_V_2.1 heterotetramerisation. (A) Wild Type K_V_6.4 co-immunoprecipitates with K_V_2.1 when co-expressed in HEK293 cells (pulling down with K_V_2.1 or HA-tagged K_V_6.4). K_V_6.4-Met419 disrupts binding to K_V_2.1, and there is significantly reduced binding of HA-tagged K_V_6.4 to K_V_2.1 when co-expressed with untagged K_V_6.4-Met419. B K_V_6.4 traffics to the plasma membrane less efficiently when co-expressed with untagged K_V_6.4-Met419, indicating a dominant negative effect. (C) Quantification of K_V_6.4 membrane localization by Pearson’s coefficient assessing colocalisation of HA and Na^+^/K^+^ ATPase membrane marker, data from three independent experiments.

We therefore conclude that the K_V_6.4-Met419 variant acts as a dominant negative subunit and significantly affects the function of K_V_6.4 (and hence in turn K_V_2.1) in the heterozygote state identified in our cohort of women who did not require analgesia during their first labour.

## Discussion

Parturition may be physiological and widely considered to be ‘natural’ but remains amongst the most painful events in life that women can experience (*34*). Labour pain is a complex experience with many biopsychosocial determinants, of which visceral nociception is fundamental and necessary. Although the cellular and molecular substrates for visceral nociception are ill defined in humans, ion channels that are important regulators of uterine sensory neurone excitability, may determine visceral nociception and hence labour pain.

Labour pain is challenging, if not impossible, to model adequately in pre-clinical laboratories. Our genetic approach in humans here was not to discover very rare Mendelian mutations that cause extreme and hence, pathological painlessness (e.g. Congenital Insensitivity to Pain). Instead, we sought to investigate SNPs that are more common, and for which frequencies in the general population are known. We hypothesised that such SNPs would be significantly over- or under-presented in a cohort of women with a less extreme, but nonetheless clinically relevant phenotype. Hence, we chose to investigate healthy nulliparous women who chose and were able to manage pain from spontaneous and uncomplicated vaginal delivery of term labour without any analgesia. In this group, there were no deficits in detection of innocuous warmth, cool or cuff-grip pressure to suggest clinically relevant sensory neuropathy. There were also no differences in cognitive test battery performance, pain-relevant personality traits or emotional function, when compared to controls. However, these women demonstrate increased pain and tolerance thresholds to a range of noxious stimuli, and significantly so for cuff-pressure pain.

Given that our painlessness phenotype is far less extreme compared to that of congenital insensitivity to pain, we did not expect that any rare-SNP(s) discovered in this study would cause a large increase in experimental pain threshold or tolerance for all stimulus modalities. Nonetheless, there is modest evidence from a study by Carvalho and colleagues that a composite of these measures obtained just before induction of labour in singleton, term pregnancies, predicts analgesic consumption, i.e. volume of local anaesthetic infused, in women who requested an epidural (*35*). The women recruited in the Carvalho cohort resembled the controls we employed in that they used analgesics during nulliparous labour. Our observation here that women who did not request any analgesic (epidural or Entonox) had an overall increase in experimental pain thresholds compared to controls is reassuringly consistent. However, the finding that cuff-pressure pain threshold was robustly and very significantly increased in those women is novel. Labour pain has visceral and somatic components, caused by contractions of uterine viscus, but also by sustained stretch or compression of pelvic floor, perineum and vagina (*4*), which occur in the later stages of labour as the foetus descends, and may be experienced as a continuous background pain on which rhythmic pain caused by uterine contractions is superimposed (*36*). Whilst speculative, the hypothesis that women with high cuff-pressure pain thresholds would report reduced intensity of continuous background pain during labour is testable.

Blinding was not feasible in our experiments and social desirability bias may explain our overall findings of increased threshold and tolerance of pain. However, such bias might be expected to also significantly lower scores for self-reported pain related traits, particularly pain catastrophizing (*11*), but that was not observed. Our data are consistent with those from other investigators who show that scores from Pain Catastrophizing Scale and Fear of Pain questionnaires do not influence self-reported or behavioural measures of labour pain (*37*). Pain is a complex experience, with sensory-discriminatory and affective-motivational aspects (*38*). We found that the test cohort had lower SFMPQ scores that pertained to the sensory, but not the affective qualities of the pain that was experienced during cold tolerance testing. In sum, we found increased threshold to pain from noxious stimuli (significantly so for cuff pressure), but no differences in cognitive, personality traits and emotional function, in women who did not require analgesics during term nulliparous labour. These findings suggest that nociceptive function is altered in these women and validate their selection to discover predisposing genetic changes in sensory neurones (nociceptors) that might influence labour pain in women: a phenotype that otherwise would confidently have been expected to be highly heterogeneous.

We detected a single SNP, rs140124801 in the gene *KCNG4* where the rare allele had a significant over-representation when compared to the general population in a cohort of 158 women who had no analgesic requirement during nulliparous labour, noting that ideally control allele frequencies would have been generated from a matched cohort of women who did require analgesia. There were 4 heterozygotes who possess the rare allele, and data on quantitative sensory and pain testing were available for 3 heterozygotes. We found that women who possess the rare allele showed a significantly increased cuff-pressure pain threshold, when compared to controls (Supplemental materials; Table S3). The rare allele of SNP rs140124801 causes a mis-sense change p.Val419Met in K_V_6.4, a silent K_V_ subunit that forms heterotetramers with K_V_2 channels and modulates their function (*23*). We, and others, show that K_V_6.4 traffics to the plasma membrane only when co-expressed with K_V_2.1 (*25*). In contrast, we found that the rare allele product K_V_6.4-Met419 failed to traffic to the plasma membrane when co-expressed with K_V_2.1. Moreover, K_V_6.4-Met419 failed to induce the hyperpolarising shift in the voltage-dependence of K_V_2.1 inactivation that is observed with K_V_6.4, likely indicating that the observed currents would be conducted by K_V_2.1 homotetrameric channels.

We have found that K_V_6.4-Met419 was unable to heterotetramerise with K_V_2.1. A possible explanation for this is gained from X-ray crystallography of K_V_2.1 homotetramer (RCSB Protein Data Bank ID: 3LNM; and see Supplemental data). Each of the four K_V_2.1 monomers contributes equally to the K^+^ ion selectivity region, which is formed by the peptide backbone carbonyl groups of the amino acids TVGYG. The side chains of Valine and Tyrosine from each of the four monomers fits within an aliphatic pocket of the adjacent monomer (composed of amino acids WWAIIS, see Figure 1C). The rare allele of SNP, rs140124801 results in Valine being changed to Methionine, the side chain of which is too large to be accommodated by this aliphatic pocket. This may be sufficient to stop K_V_6.4 forming a heterotetramer with K_V_2.1 and would be predicted to disrupt the close packing of the peptide backbone carbonyl groups of the ion selectivity region.

For K_V_6.4 to modulate labour pain it needs to be expressed in an appropriate part of the sensory nervous system. We focused on uterine sensory neurones, but this does not negate the possibility that K_V_6.4 also exerts influence elsewhere in the nervous system, *KCNG4* mRNA also being expressed in regions of the spinal cord and brain (Figure S7). We observed K_V_6.4 expression in Trpv1 and Na_V_1.8-positive mouse uterine sensory neurones, consistent with the observation that sensory neurones innervating deep tissues display comparatively high Trpv1 expression (*39*). Results from unbiased single-cell RNA-sequencing of mouse DRG obtained from cervical to lumbar levels reveal no specific coexpression of K_V_6.4 in nociceptive Trpv1/Scn10a expressing neurones (*32*). However, single-cell RNA-sequencing of colonic sensory neurones identified that K_V_6.4 does co-localise with Trpv1 and Na_V_1.8 (*29*), consistent with our findings here that K_V_6.4, Trpv1 and Na_V_1.8 are coexpressed in uterine sensory neurones from T12-L2 and L5-S2 DRG. Taken together, these data suggest that K_V_6.4 might be a marker for sensory neurones that innervate the viscera. Due to the restricted expression of *Kcng4* in a particular sensory neurone type, expression of K_V_6.4-Met419 is expected to reduce excitability specifically for this class of sensory neurones.

For the rare allele rs140124801 to modulate labour pain it needs to cause a significant change in K_V_6.4-influenced neuronal activity, and to do so in the heterozygote state. Our electrophysiology and cell trafficking studies showed that the mutant K_V_6.4-Met419, as opposed to K_V_6.4, had no effect on K_V_2.1 function, nor was it trafficked to the plasma membrane. Transfection of K_V_6.4 into mouse sensory neurones produced a more hyperpolarised voltage-dependence of inactivation for the predicted heterotetrameric K_V_2/silent K_V_ channel component of *I*_K_ than when K_V_6.4-Met419 was transfected, further supporting the hypothesis that the loss-of-function K_V_6.4-Met419 results in more K_V_2.1 activity at positive voltages. K_V_2.1 is known to contribute to the after-hyperpolarisation duration, intra-action potential refractory period, and thus regulate neuronal excitability (*27*). Hence, we anticipated that a K_V_6.4-Met419-induced deficit in K_V_2.1 function would likely result in fewer action potentials and thus less pain during periods of sustained nociceptor activity, such as that occurring with uterine contractions during labour. Although we did not observe a difference in the after-hyperpolarisation duration or action potential frequency between sensory neurones transfected with K_V_6.4 or K_V_6.4-Met419 (possibly due to the continual current injection used), we did find that a larger amount of current was required to cause K_V_6.4-Met419 expressing neurones to fire actions potentials and thus conclude that the mutation confers reduced neuronal excitability. Critically, we observed that K_V_6.4-Met419 has a dominant negative effect on K_V_6.4, regarding modulation of the voltage-dependence of inactivation for K_V_2.1. This result likely explains the reduction in labour pain seen in individuals in our cohort who were heterozygotes for the SNP rs140124801 rare allele.

There is a growing understanding of the distinctions between the neural pathways for pain from visceral and somatic tissues: each have evolved nociceptors that sense damage in different physical environments (*40*). Our findings suggest a key role for K_V_6.4 in specifically regulating nociceptor excitability, and hence pain, in normal labour. K_v_6.4 is also expressed in other parts of the nervous system (Figure S7) and its expression in non-neural tissues is unknown. However, we found that women carrying the rare allele *KCNG4* managed nulliparous labour without analgesics, have higher experimental pain thresholds but are otherwise healthy without any psychological or cognitive abnormalities. Their phenotype suggests that the loss of modulatory effects of K_V_6.4 is non-pathogenic in other parts of the nervous system and non-neural tissues. If druggable, K_v_6.4 would be a potential target for modulating labour pain without the maternal and neonatal side effects inherent in other analgesic interventions in this setting. Our data also raise the question of whether K_V_6.4 has roles in other painful visceral disorders, both within and outside the female genital tract. One closely related context would be primary dysmenorrhea, which is characterised by severe pain associated with uterine contraction during menstruation (*41*). Further development of selective K_V_6.4 pharmacological agents is required to fully probe the role of K_V_6.4 in visceral pain.

## Materials and Methods

For detailed Materials and Methods, see Supplementary Materials

### Study Design

The purpose of this study was to identify possible genetic contributions affecting the levels of pain experienced by women during their first labour. We screened >1000 cases and ascertained a cohort of 189 healthy women who had given birth without any use or request for analgesics. A DNA sample was collected from each (158 provided DNA samples of sufficient quality for further analysis). 39 of this cohort consented for comprehensive psychometric and quantitative sensory testing (QST) and were compared to 33 matched controls who used analgesics during labour and delivery of their firstborn. Psychometric evaluation focused on emotional function, cognitive abilities and personality traits that may influence the experience of pain. QST comprised ascertaining somatic sensory detection, pain and tolerance thresholds of thermal and mechanical stimuli in controlled experimental settings.

The chronologically first 100 DNA samples collected were designated a discovery cohort and underwent full exome sequencing. We mined the data for rare allele SNPs that were statistically under or over represented (by Chi-squared test with two tails and Yates correction) in the discovery cohort after false discovery rate (FDR) correction and identified a p.Val419Met alteration in *KCNG4*, which was then sequenced by Sanger Sequencing methodology in the 58 women replication cohort. This data led us to investigate the potential contribution of the *KCNG4* encoded protein K_V_6.4 to the phenotype observed.

K_V_6.4 was cloned, tagged and the p.Val419Met introduced by site-directed-mutagenesis with pathogenicity assessed by electrophysiological characterization and assessment of intracellular localization in HEK293 cells. The finding of the alteration behaving as a dominant negative, loss of function mutation led us to investigate the physiological role of K_V_6.4 in uterine sensory neurones. Using retrograde labelling we identified uterine sensory neurones (89 were collected) and performed single-cell qRT-PCR to identify K_V_6.4 expression alongside binding partner K_V_2.1 and nociceptor marker genes. Finally, having identified K_V_6.4 expression alongside K_V_2.1 in a significant proportion of uterine sensory neurones, we further characterized the effects of expressing the K_V_6.4 minor allele in mouse sensory neurones by whole-cell patch-clamp recording, confirming our initial findings of K_V_6.4 Met419 as a loss of function mutation and demonstrating decreased excitability of neurones transfected with K_V_6.4-Met419.

### Statistical Analyses

For psychometric and quantitative sensory testing, statistical analyses were performed with R Studio (Version 1.1.442). The mean and standard deviation were generated for each outcome variable for test and control cohorts. Shapiro-Wilk tests and F-tests were used assess data normality and differences in variances. Differences between the means of each outcome variable in test and control cohorts were assessed using tests for two independent samples, using Student’s *t*-test, Welch’s *t*-test or Mann-Whitney U tests when the relevant assumptions were met. The level of statistical significance was adjusted using Sidak’s correction. The correction applied to multiple outcomes associated with each domain of assessments: questionnaires, CANTAB, sensory detection, pain thresholds and tolerance.

For statistical assessment of the genetic data collected in this study, enrichment of amino acid altering SNPs was assessed by exome sequencing, with exome vcf, bam and bam.bai files iteratively analyzed to extract data on all SNPs in or near to exons, including the depth and quality of the sequence data, and the alleles detected (*9*). For each SNP the allele frequencies were compared to normal values derived from the 1000 genomes project and exome variant server, and deviations assessed for significance using a Chi-squared test with two tails and Yates correction. The resulting *P* values were subject separately to a Bonferroni and FDR correction, as approximately 100,000 SNPs were assessed in our fSNPd method.

We then focused only on ion channels, defined as being identified by the Gene Ontology Term GO:0005216 (423 genes, which were also hand curated and checked against a Pfizer/Neusentis database that had been shared with us). There were 28 SNPs found in ion channels and each was in a different gene; there was only one detected SNP in *KCNG4*. Eight of these SNPs were then eliminated because the protein change caused by the rare allele was common in the orthologous mammalian proteins. For the remaining 20 SNPs, we determined the allele frequency of each of the by use of the Integrated Genome Viewer examining the cohort’s exome bam files individually. This led to the elimination of 19 SNPs, 14 as the common allele frequency was 100% and program errors in assigning alleles within our discovery cohort had falsely appeared to show a deviation from 100%, and five because of misalignment of reads to homologous genes leading to errors in SNP allele calling and SNP allele frequency calculation.

Further statistical tests used to assess differences between K_V_6.4 and K_V_6.4-Met419 throughout the study are unpaired *t*-tests and ANOVA with Bonferroni’s multiple comparison post-hoc test, as described in the relevant figure legends and Supplementary Methods. Differences between groups were considered significant at a *P* value < 0.05, and were tested using GraphPad (Prism5.0, California, USA).

## Acknowledgments

The authors would like to thank the mothers who participated in this study. We acknowledge the support from the National Institute of Health Research who facilitated identification of potential participants. We thank midwives and other clinicians who assisted with case ascertainment and invited potential participants on our behalf. We would like to acknowledge Ingrid Scholtes and staff at the Cambridge NIHR Clinical Investigation Ward who cared for our participants during their visit.

## Funding

MCL, DKM, DW, and CGW acknowledge funding from Addenbrooke’s Charitable Trust and the NIHR Cambridge Biomedical Research Centre. MN was funded by the Wellcome Trust (200183/Z/15/Z); JH and ESS by a Rosetrees Postdoctoral Grant (A1296) and the BBSRC (BB/R006210/1); GC and ESS by Versus Arthritis Grants (RG21973); VBL and FR by the Wellcome Trust (106262/Z/14/Z and 106263/Z/14/Z) and a joint MRC programme within the Metabolic Diseases Unit (MRC_MC_UU_12012/3). EF, GI and CB were funded by the Cambridge NIHR Biomedical Research Centre Integrative Genomics theme and LAP by a BBSRC-funded studentship (BB/M011194/1). We thank Professor Naomichi Matsumoto and Professor Hirotomo Saitsu for the kind use of their KCNG4 vector. We would like to thank Walid Khaled for the kind loan of the Nucleofector key to electrophysiological experiments.

## Author contributions

MCL, MSN, FR, DW, DKM, ESS and CGW made substantial contributions to the conception and design of this work. MCL, GI, CB and CGW made substantial contributions to the acquisition of clinical data. MSN, JRFH, VBL, LAP, GC and PE made substantial contributions to the acquisition of cell and molecular biology data. MCL, MSN, JRFH, VBL, LAP, ID, GC, KS, FR, EVF, PE, ESS and CGW made substantial contributions to the analysis and interpretation of data for the work. All authors were responsible for the drafting of the work or revising and giving final approval of the version to be published, and agree to be accountable for all aspects of the work.

## Material requests correspondence

ESS or CGW

