## Supplemental materials for "Human labour pain is influenced by the voltage-gated potassium channel K_V_6.4 subunit"

#### METHODS

##### Case ascertainment and recruitment

Labour pain is a complex experience and difficult to quantify (*1*). Epidurals and inhalational analgesia are currently the most effective forms of pain relief in labour (*2*). Hence the rate of epidural use is a recognized surrogate measure for pain in clinical trials that assess the effectiveness of other forms of analgesia in labour (*3*). The use of inhalational analgesia is far commoner, particularly in nulliparous parturients where labour is considered more painful. A UK survey suggests that Entonox® use in labour at 80% and first-time mothers were more likely to use labour analgesia (*4*). Hence, the phenotype for less painful labour was defined operationally as nulliparous parturients who did not request nor use epidural, inhalational or opioid-based analgesia. This behavioural definition would have captured individuals with *SCN9A* channelopathy who reported entirely painless labour (*5*).

The studies commenced in October 2012 after National Research Ethics Service and Human Research Authority approval (Reference: 12-EE-0369) was granted. For the first study (Study A), potential participants were identified based at maternity units in the United Kingdom and invited via post to contact the research team. The post included an information sheet stating that the study sought "to use genetic analysis to look for variations in genes in women who do not feel as much pain as might be expected during childbirth, and determine whether such variation in pain experience might be related to genetic differences' and the invitation was for women that 'have had a baby and according to our records, you required minimal or no pain relief during the birth of your first child".

All potential participants who contacted the research team were further screened via telephone interview for eligibility (Supplemental Table S1). Eligible participants were posted study information and a saliva sampling kit (Oragene®-DNA, OG-500, Genotek), with a self-addressed return envelope. Participants did not receive any financial incentive for the genetic study.

In the second study (Study B), women who had consented to the genetic study and for whom exome sequencing was successful were invited to the Cambridge NIHR Clinical Investigation Ward for further study. Those who were eligible (Supplemental Table S1) and consented to participate comprised the test cohort.

Women who met the study criteria but who required analgesia during labour served as case controls. Controls were informed via participant information leaflet we have *"identified women who did not use painkillers during the birth of their first child. However, we are still unsure whether they are actually less sensitive to pain. In order to find out, we need to test their pain sensitivity and compare their results to women who did used an Epidural or Entonox (gas and air) for pain relief during their first labour"*. Controls were selected to match age at delivery of first-born, location of maternity unit and age at study visit. A total of 1029 invitations were sent by post. Where available, data on birth weight of baby and head circumference were recorded.

Participants were reimbursed up to maximum of 25GDP for time in addition to travel expenses for the two-hour visit. All participants and the researchers who communicated directly with them remained blind to genotype during the study.

A single research assistant in the same temperature-controlled room conducted all assessments. Participants were seated for the assessment and rest breaks were provided between assessments to minimize fatigue. Instructions for each assessment were read from a written script. These assessments were completed in the following sequence: (1) questionnaires administered on paper (HADS, PCS, MHLC and LOTR), (2) quantitative sensory testing (QST) to determine stimulus detection, pain and tolerance thresholds and (3) computerized cognitive assessments implemented on (CANTAB®) and

#### Cambridge Neuropsychological Test Automated Battery (CANTAB)

The cognitive assessments were drawn from the Cambridge Neuropsychological Test Automated Battery (CANTAB) (http://www.cambridgecognition.com/). The computerised tests required finger-tap responses via touchscreen and are largely independent of verbal instruction. CANTAB software was deployed on an XGA-touch panel 12-inch monitor (Paceblade Slim-book P120; PaceBlade Technology). The sequence of tasks employed in the study was as follows: Motor Screening Task (MOT), Spatial Working Memory (SWM), Rapid Visual Information Processing (RVIP), Intra- Extra-Dimensional Set Shift (IED) and One-Touch Stockings of Cambridge (OTS). Descriptions of each task are provided below. All tasks were performed using the index finger of the dominant hand

**Motor Screening Task (MOT)**

Coloured crosses are presented in different locations on the screen, one at a time. The participant must select the cross on the screen as quickly and accurately as possible. Outcome measures are (a) mean latency and (b) mean error, which reflect accuracy.

#### Spatial Working Memory (SWM)

The task assesses ability to retain spatial information and manipulate items in working memory. It is considered a sensitive measure of frontal lobe and executive dysfunction. This is a self-ordered task which also assesses heuristic strategy.  Several coloured squares (box) are displayed in random locations on the touch screen. There is pre-set number of boxes with a blue token. The participant taps on a box to uncover a blue 'token' and place that token into a 'bin'. The participant must remember which box has been tapped or emptied. The number of boxes is gradually increased until the participants is searching for tokens in a total of eight boxes. The colour and position of boxes used are changed from trial to trial to discourage use of stereotyped search strategies. Outcome measures are (a) strategy, for which the fully efficient strategy would result in no boxes being revisited. A high score represents poor use of this strategy and a low score equates to effective use, and (b) total errors, which is the number of times a box is selected that cannot contain a blue token and therefore should not have been visited by the subject.

#### Rapid Visual Information Processing (RVP)

A white box is shown in the centre of the screen, inside which digits from 2 to 9 appear in a pseudo-random order, at the rate of 100 digits per minute. Participants are asked to detect target sequences of digits (for example, 2-4-6, 3-5-7, 4-6-8) and respond by tapping on a button-box as quickly as possible. Outcome measures are (a) sensitivity index A', which reflects how good the subject is at detecting target sequences, regardless of error tendency A score close to +1.00 indicates that a high true positive rate, and (b) response criterion B', which reflects the tendency to respond regardless of whether the target sequence is present. A score close to +1.00 indicates a high true negative rate.

#### Intra-Extra Dimensional Set

This task assesses visual discrimination and attentional set formation maintenance, shifting and flexibility of attention. IED task requires participants to learn the rule and select the correct icon (a specific shape or line). The task builds in complexity as distractors are added and the rule changes. The rule changes are both intra-dimensional (e.g. shapes are still the relevant set, but a different shape is now correct) and extra-dimensional (e.g. shapes are no longer the relevant set, instead one of the line stimuli is now correct). Outcome measures are (a) total errors (adjusted), which is a measure of the participant's efficiency. Whilst she may pass all nine stages, a substantial number of errors may be made in doing so. The errors are adjusted to account for those who fail at any stage of the test and hence have had less opportunity to make errors, (b) number of stages completed, and

(c) total trials (adjusted), which is the number of trials completed on all attempted stages for each stage not attempted due to failure at an earlier stage.

**One Touch Stockings of Cambridge**

This task is a test of executive function, based upon the Tower of Hanoi. The participant is shown two displays containing three coloured balls. The displays are presented in such a way that they can be easily perceived as stacks of coloured balls held in stockings suspended from a beam. The participant is shown how to move the balls in the lower display to copy the pattern in the upper display and completes one demonstration problem, where the solution requires one move. The participant must then complete three further problems, one each requiring two moves, three moves and four moves. Next the participant is shown further problems and must work out mentally the number of moves the solutions require and then select the appropriate box at the bottom of the screen to indicate their response. Outcome measures are (a) mean choice to correct, which is the mean number of attempts to the correct response, and (b) mean latency to correct, which is the overall latency (time required) to the correct response.

#### Quantitative sensory testing

Stimulus detection and pain thresholds for heat and cold were determined by applying a 3x3 $cm^2$ thermode on the volar surface of the non-dominant mid-forearm (TSA, Medoc, Israel). The procedure was adapted from a clinical research protocol(*6*), for which the research assistant received formal training (Universitsmedizin Mannheim). Stimulus detection and pain thresholds were determined using increasing or decreasing temperature ramp of 1°C.$s^{-1}$ from a baseline temperature of 32°C, with low and high safety cut-offs at 0 and 50°C respectively. Participants were instructed to click on a button when they first experience the required sensations. Four trials each with an inter-trial interval of 10s were employed to assess heat and cold stimulus detection thresholds. Three trials with a longer inter-trial interval of 30s were employed for heat and cold pain thresholds to minimize risks of burn.

Pressure detection and pain threshold were determined by cuff algometry(*7*) applied to the dominant upper arm. The circumference of the upper arm was measured to determine the appropriate sphygmomanometer cuff size. A digital metronome (Korg MA-1, UK) was used to guide manual inflation of the cuff at 10 mmHg every 5s. The participant was instructed to verbally report when the point the cuff was felt to be ‘just gripping’ and when the gripping became just about painful, at which point the cuff pressure was rapidly released. The pressures at thresholds were recorded. The participant was then asked to indicate when all evoked sensation in the arm had resolved. The entire procedure was repeated thrice.

Pain tolerance was measured as latency to withdrawal from immersion of hand in a cold water bath (3 °C) (*8*). The participant was instructed to immerse her non-dominant hand and wrist into a circulating cold-water bath (RW2025G, Medline Scientific UK) and to withdraw the hand *ad libitum* when pain became intolerable. The maximum duration of cold-water immersion allowed was 180s, after which the participant was told to remove her hand from the water bath. All participants were told that there was a maximum allowable duration for immersion for safety but not the exact duration to avoid anticipatory effects.

The skin temperature of the hand dorsum was measured (NC 150, Microlife, Switerland) within 60s pre-immersion and 10s post-immersion (after the hand was wiped dry). Participants were then asked to rate peak intensity of pain during immersion using a 100mm visual analogue scale (VAS) with the left and right anchors labelled as ‘no pain’ and ‘worst imaginable pain’ respectively. They were also asked to estimate when the intensity of pain peaked during the period of immersion (100mm VAS; 0mm and 100mm represented the times of hand immersion and hand withdrawal respectively). Finally, participants completed the Short-Form McGill Pain Questionnaire (SQ-MPQ). The questionnaire comprises 15 pain descriptors: 11 pertain to sensory-discriminatory aspects (e.g. ‘hot-burning’), and the rest pertained to affective-motivational (e.g. ‘cruel-punishing’) of the pain experienced during cold-water immersion of the hand (*9*).

#### Genetic analysis of non-synonymous functional single nucleotide polymorphism alleles

For the genetic analysis of the discovery cohort we used the fSNPd approach (*10*). In brief, the hypothesis is that some individuals with a defined phenotype (in our case reduced labour pain inferred by the absence of analgesia requirement during labour) could have genetic predisposition(s) that explain their difference in phenotype. To be detected, such a genetic predisposition would have to be dominantly inherited and often penetrant: this is the case with many known autosomal dominant Mendelian genetic disorders such as Tuberose Sclerosis and Neurofibromatosis type 1 where the phenotype is variable (and can be incomplete) despite an individual carrying the known pathogenic familial mutation. The fSNPd approach further hypothesizes that the phenotype will not be caused by very rare genetic mutations, but by the rare alleles of known SNPs where the allele difference is protein changing. Examples of such SNPs exist that only cause a human phenotype when the heterozygous individual is exposed to a specific environmental insult or trigger, e.g. aminoglycoside induced deafness (*11*) and SNPs rs267606617 and rs267606618; and carbamazepine associated toxic epidermal necrolysis and rs3909184 (*12*).

An exome analysis was performed on the genomic DNA of the 100 individuals of the discovery cohort by Beijing Genomics Institute using the Agilent 51M kit sequenced to an average of 50-fold coverage, as previously described (*13*). Such an analysis does not include all coding exons of all human genes, and for this reason SNPs in some genes are not assessed, and other SNPs were not assessed in all individuals in the discovery cohort (*10*). The exome vcf, bam and bam.bai files were iteratively analysed extracting data on all SNP in or near to exons, including the depth and quality of the sequence data, and the alleles detected (*10*). For each SNP the allele frequencies were compared to normal values, and deviations assessed for significance using a Chi-squared test with two tails and Yates correction. The resulting *P* values were subject separately to a Bonferroni correction and false discovery rate (FDR) correction, as approximately 100,000 SNPs were assessed in our fSNPd method. We then filtered only for clear-cut protein changing SNPs (mis-sense mutations predicted deleterious by SIFT, nonsense mutations, splice site mutations, start codon mutations, and within-exon deletions and duplications), as such changes are potentially more amenable to function tests of pathogenicity; reducing from 18,106 SNPs prior to SIFT and pathogenicity analysis to 3,596 afterwards. We then further filtered only for SNPs within ion channel genes, as members of this group of genes have already been implicated in Mendelian pain disorders, and testing techniques for ion channel function are well established; resulting in 28 SNPs (*14*). For all SNPs, especially those whose rare allele frequency is < 5 %, geographical and ethnic differences must be considered; rs140124801 has a rare allele frequency in EVS of 0.0051 (cohort size 6500), in gNOMAD Europeans = 0.0072 (cohort size 18,878), 1000 Genome = 0.0048 (cohort size 2,504), and our population were Caucasian and predominantly born in the United Kingdom.

In the discovery cohort we assessed all individual bam files with the Integrated Genome Viewer to determine which rs140124801 alleles were present, blind of the fSNPd results. All individuals predicted to have the rs140124801 rare allele were Sanger sequenced and complete concordance was found. Primers were designed with Primer3 and are available on request. Genomic DNA of the 58 individuals in the replication cohort was Sanger sequenced to determine the SNP rs140124801 allele frequencies. The allele frequency and number of heterozygotes of rs140124801 was assessed in combined cohort of discovery and replication by $Chi^2$ test with two tailed and Yates correction (for small numbers), using the more conservative control population allele frequency of 0.0072 for heterozygote carriers.

We assessed the effects on protein sequence and protein function of the *KCNG4* SNP rs140124801 alleles by use of bioinformatic resources within the Human Genome Browser, NCBI BLASTP for protein sequence comparisons and Conserved Domains (CD search) for detecting if the amino acid change occurred within a known protein domain and SIFT for pathogenicity prediction.

#### Modelling of *KCNG4* alleles on the $K_v2.1/K_v6.4$ heterotetramer

We used the X-ray crystallography derived structure of rat $K_V2.1$ homotetramer (RCSB Protein Data Bank ID: 3LNM) to model the effects of the rare allele of rs140124801 (*15*). Rat and human subunits form both $K_V2.1$ homotetramers and 3:1 $K_V2.1$:$K_V6.4$ heterotetramers. Rat was the closest species to humans with a published $K_V2.1$ protein structure. Rat and human $K_V2.1$ proteins are 94% identical and 79% identical for $K_V6.4$. However, restricting the alignment to the 78 amino-acid region physically adjacent to the $K^+$ selectivity filter (the pore loop from transmembrane region 5 to transmembrane region 6, which includes the $K^+$ selectivity filter) rat and human $K_V2.1$, they are identical and rat and human $K_V6.4$ is 96% identical (with no amino acid changes in the aliphatic pocket or selectivity filter).

We used structure 3LNM and the Cn3D software (*16*) to examine the $K^+$ selectivity filter of the $K_V2.1$ homotetramer to look at the sites of interaction of each of the four individual $K_V2.1$ proteins, and produced images where proteins and individual amino acids were identifiable, or could be omitted from the whole tetrameric structure.

The $K^+$ ion selectivity filter is formed by the peptide backbone carbonyl groups of the amino acids TVGYG of each of the four $K_V$ subunits. This forms a narrow central channel through which potassium ions ($K^+$) can flow out of the cell. Each $K_V$ subunit forms an identical quarter of the tetramer structure about the selectivity filter region central pore. The $K_V2.1$ homotetramer model reveals the side chains of the Valine and Tyrosine of each subunit selectivity filter TVGYG protruding into a highly conserved "aliphatic pocket" (with canonical sequence WWAIIS, see Figure 1C) in the adjacent $K_V$ subunit. In this model the Valine-419 of $K_V6.4$ can be accommodated identically compared with the equivalent Valine-374 of $K_V2.1$. However, the larger aliphatic side chain of 419-Methionine in the $K_V6.4$ SNP would not be able to be accommodated within the aliphatic pocket, and hence would disrupt the ion selectivity region of the $K_V2.1/2.1/2.1/6.4$ heterotetramer.

#### DNA constructs and antibodies

A full-length human *KCNG4* cDNA clone was purchased from Source bioscience (IRCMp5012B0629D) and cloned in-house into a pcDNA3 based expression plasmid (CMVgenex-polioIRESmCherry) both with or without a C-terminal HA tag. The p.Val419Met mutation was introduced by site-directed mutagenesis (Stratagene) according to the manufacturer's protocols and sequences of the plasmids were confirmed by Sanger Sequencing. The clone expressing $K_V2.1$ alongside a nuclear GFP reporter in the pCAGGS-IRES2-nucEGFP vector has been described previously(*17*).

Antibodies used were anti-HA mouse monoclonal (12B12, #MMS-101P, Biolegend), anti-$Na^+/K^+$ ATPase rabbit monoclonal (ab76020, Abcam), anti-mCherry rat monoclonal (M11217), anti-β-actin mouse monoclonal (a2228, Sigma), anti-$K_V2.1$ rabbit polyclonal (APC-012, Alomone), anti-$K_V6.4$ mouse monoclonal (N458/10, NeuroMab), anti-$K_V2.1$ mouse monoclonal (K89/34, ab192761, Abcam), and anti-HA rabbit monoclonal (C29F4 #3724, Cell Signalling).

#### Immunofluorescence analysis and confocal microscopy

HEK293 and HeLa cells were cultured on poly-L-lysine coated coverslips and transfected as described above. 48-hours after transfection, cells were fixed by 10 minutes incubation in 4 % paraformaldehyde. Cells were permeabilized by 10 minutes incubation in 0.3 % Triton-X100 solution followed by 30 minutes at room temperature in 5 % BSA solution. Alternatively, when staining for $Na^+/K^+$ ATPase, cells were fixed and permeabilized by emersion in ice cold methanol. Fixed cells were then stained with primary antibodies for 1 hour in 5 % BSA and fluorescent secondary antibody also for 1 hour. Secondary antibodies used were Alexa Fluor 488 donkey anti-mouse, Alexa Fluor 546 goat anti-rabbit, Alexa Fluor 546 donkey anti-mouse, Alexa Fluor 633 goat anti-mouse (all from Life Technologies). Coverslips were mounted onto glass slides using Prolong Diamond Antifade Mountant with DAPI (Molecular Probes). Cells were visualised with an LSM880 confocal microscope.

#### Co-immunoprecipitation

HEK293 cells were transfected with $K_V2.1$ and $K_V6.4$ plasmid constructs as described in the associated figures, and harvested after 3 days. Co-immunoprecipitation was carried out using the Dynabeads Co-Immuniprecipitation Kit (Life Technologies) according to the manufacturers protocols. Antibodies used were anti-HA mouse monoclonal (12B12, #MMS-101P, Biolegend), anti-$K_V2.1$ rabbit polyclonal (APC-012, Alomone), anti $K_V2.1$ mouse monoclonal (K89/34, ab192761, Abcam), and anti-HA rabbit monoclonal (C29F4 #3724, Cell Signalling). IP buffer supplied in the kit was supplemented with 80mM NaCl.

#### Cell lines and culture conditions

HEK293 and HeLa cells were cultured in 90 % Dulbecco's modified Eagle medium (DMEM) supplemented with 10 % fetal bovine serum (FBS), 100 U/ml penicillin-streptomycin (pen-strep), and 2 mM L-glutamine at 37 °C, 5 % $CO_2$, 100 % humidity. Transfections were carried out using FugeneHD transfection reagent (Promega) according to the manufacturer's protocols. For co-expression studies, cloned $K_V$2.1 and $K_V$6.4 constructs were transfected at a ratio of 1:2. Cells for experiments were plated out on glass coverslips for immunostaining or 35 mm plastic dishes for electrophysiological recordings, 1-2 days prior to the experiment.

#### Electrophysiological characterization of *KCNG4* SNP rs140124801 alleles and *KCNB1* in HEK293 cells

Whole-cell recordings from transfected HEK293 cells were performed using 1-2.5 MΩ resistance fire-polished borosilicate glass electrodes filled with an internal pipette solution containing (in mM): KCl (110), $K_4$-BAPTA (5), HEPES (10), $MgCl_2$ (1), $K_2$ATP (5), pH 7.3, 281 mOsm/kg. Cells were continuously superfused with bath solution containing (in mM): NaCl (145), KCl (4), HEPES (10), D-glucose (10), $CaCl_2$ (1.8) $MgCl_2$ (1), pH 7.4, 300 mOsm/kg, at room temperature (20-24 °C). Potassium currents ($I_K$) were recorded in voltage clamp mode using an Axopatch 200B connected through a Digidata 1440A A/D converter and pCLAMP software (version 10, Axon Instruments). The calculated linear leakage current was digitally subtracted offline for all current measurements. Potassium currents were elicited by step depolarisations from a holding potential of −90 mV to various test potentials. The voltage-dependence of activation was determined from tail currents recorded from a 200 ms voltage step to −60 mV following these various test potentials. The normalised tail currents were plotted against the voltage of the step depolarisations and fit with a Boltzmann function. A double pulse protocol was used to measure the voltage-dependence of steady-state inactivation. The protocol consisted of a 5 s prepulse that ranged between −110 to +40 mV from a holding potential of −90 mV followed by a 200 ms test pulse to +50 mV. Normalised currents during this test pulse were plotted against the prepulse voltage and fit with a Boltzmann function. The time course of recovery from inactivation was investigated by applying a 5 s prepulse to +20 mV from a holding potential of −90 mV and applying a 200 ms test pulse to +20 mV at various time intervals after the conditioning prepulse. Recoveries from inactivation time courses were fit to a single exponential function.

#### Animals

Adult C57BL/6J mice (Envigo) were conventionally housed in groups of 4-5 with nesting material and a red plastic shelter and various enrichment toys; the holding room was temperature controlled (21 °C) and mice were on a 12-hour/light dark cycle with food and water available *ad libitum.* Work was regulated under the Animals (Scientific Procedures) Act 1986 Amendment Regulations 2012 following ethical review by the University of Cambridge Animal Welfare and Ethical Review Body.

#### Single-cell qRT-PCR of mouse uterus innervating sensory neurons

Uterus innervating sensory neurons were retrograde labelled and the mRNA transcript expression for genes of interest determined using methodology previously described for other visceral organs (*18-20*). Female C57BL/6J mice (10-12 weeks) were used. Following laparotomy, 2 injections (~2.5 µl/injection) of Fast Blue (FB: 2% in saline) were made, one into each uterine horn adjacent to the cervix. Following recovery, animals were provided a soft, glucose-enriched diet and prophylactic post-operative analgesia (buprenorphine 0.05-0.1 mg $kg^{-1}$). After 5-8 days, mice were killed by cervical dislocation and two primary cultures made from DRG T12-L2 (TL) and L5-S2 (LS), respectively. Dissected DRG were incubated in Lebovitz L-15 Glutamax (Thermo Fisher Scientific, UK) media containing 6 mg $ml^{-1}$ bovine serum albumin (BSA, Sigma-Aldrich) and 1 mg $ml^{-1}$ collagenase type 1A (Sigma-Aldrich, UK) for 15 min at 37 ºC in 5 % $CO_2$. After a further 30 min incubation in L-15 media containing 1mg $ml^{-1}$ trypsin (Sigma-Aldrich) and 6 mg $ml^{-1}$ BSA, ganglia were triturated and dissociated cell-containing supernatant collected by repeat brief centrifugation (5 x 500 *g*). TL and LS neurons were plated on poly-D-lysine coated coverslips (BD Biosciences, UK) and incubated in L-15 growth media (containing 2 % penicillin/streptomycin, 24 mM $NaHCO_3$, 38 mM glucose and 10 % fetal bovine serum). Fluorescently labelled FB-positive cells were picked manually by pulled glass pipette into 9 µl mastermix (containing 5 µl CellsDirect 2 x reaction buffer (Invitrogen, UK), 2.5 µl 0.2 x primer-probe mix against genes of interest, 0.1 µl SUPERase-in (Ambion, USA), 1.2 µl TE buffer (Applichem, Germany) and 0.2 µl Superscript III Reverse Transcriptase-Platinum Taq mix (Invitrogen, UK)), bath samples were collected as negative controls and all samples immediately frozen on dry ice. Prior to 1:5 dilution in TE buffer, reverse transcription and preamplification of cDNA was performed by thermal cycling (50 °C for 30 min, 95 °C for 2 min, then 24 cycles of 95 °C for 15 s, 60 °C for 4 min). Gene-specific Taqman qPCR assays were then run (Taqman Assay ID: *Kcng4*, Mm01240890_m1; *Kcnb1*, Mm00492791_m1; *Trpv1*, Mm01246300_m1; *Scn10a*, Mm00501467_m1; *Gapdh*, Mm99999915_g1; Applied Biosystems) with the following thermal cycling protocol (50 °C for 2 min, 95 °C for 10 min, then 40 cycles of 95 °C for 15 s, 60 °C for 1 min). The expression of glyceraldehyde-3-phosphate dehydrogenase (*Gapdh*) acted as an internal positive control and was present in all single-cell RT-PCR products and absent in bath control samples. 15 uterine sensory neurons per region (TL and LS) per mouse ($N$ = 3) were collected. In total, 90 neurons were collected, and photos taken for analysis of cell diameter. qPCR products were detected in 89 neurons and quantitative assessment of gene expression was determined by quantification cycle values less than the threshold of 35 considered positive.

#### Whole-cell patch-clamp recordings

Primary DRG cultures from C57BL/6J mice (8-10 weeks) were generated using the methodology described for single-cell qRT-PCR experiments with the following exceptions. From each mouse, DRG T10-S1 were dissected and pooled into a single primary culture. After trituration, in order to purify the DRG culture to improve transfection efficiency, dissociated cells were subjected to a 3.5 % BSA (in L-15 media) density gradient centrifugation (20 mins at 20 *g*) and the supernatant discarded. The remaining purified dissociated neurons were washed once in L-15 growth media before resuspension in 100 µl of Mouse Neuron Nucleofector solution (Amaxa Mouse Neuron Nucleofector Kit, Lonza, UK) containing 4.5 µg plasmid of either wild-type $K_V$6.4 or $K_V$6.4-Met419 in a CMV-KCNG4-polioIRESmCherry expression cassette. Incorporation of the plasmid was achieved by electroporation (Program O-0005; Nucleofector IIb, Lonza, UK) and cells plated on poly-D-lysine/laminin coated coverslips (BD Biosciences, UK) and incubated at 37 °C in 5 % $CO_2$ and L-15 growth media. Electrophysiology experiments were conducted 48-hours post-transfection, neurons positive for mCherry fluorescence were selected following excitation with a 572 nm LED (Cairn Research, UK).

To assess voltage-gated $K^+$ currents, patch pipettes were pulled using a P-97 pipette puller (Sutter Instruments, USA) with typical resistances of 3-5 MΩ and back-filled with the pipette solution containing (in mM): KAspartate (110), KCl (30), $MgCl_2$ (2), $CaCl_2$ (1), $Na_2ATP$ (5), EGTA (2), cAMP (0.1), HEPES (10), pH 7.4. Recordings were obtained using a Multiclamp 700A amplifier (Molecular Devices, USA) in the voltage-clamp mode and digitised using a Digidata 1440A (Molecular Devices). Voltage errors were minimized using 70% series resistance compensation. Mouse neurons were continuously superfused with the bath solution containing (in mM): N-methyl-D-glucamine (NMDG; 140), KCl (5), $MgCl_2$ (1), $CaCl_2$ (1.8), glucose (10), HEPES (5), pH 7.4. The osmolality of both solutions was adjusted to 300-310 mOsm. Cells with series resistance values greater than 15 MΩ were omitted from analysis.

$I_K$ activation and inactivation protocols (Figures 3 and S5) were applied after achieving whole-cell rupture. Using a rapid change perfusion system (Intracel EVH-9, UK), 100 nM Stromatoxin-1 (Alomone, Israel) in bath solution was applied to the cells for 3 minutes prior to repeating activation and inactivation protocols. Thus ScTx-sensitive $I_K$ was determined by subtraction of the post-ScTx $I_K$ from the pre-ScTx $I_K$ in pClamp software (Molecular Devices). The voltage-dependence of $I_K$

activation was fitted with the following Boltzmann equation: $y = t / (1 + \exp((V_{50} - E)/k))$, where E is the applied voltage, $V_{50}$ is the voltage at which 50 % of the channels are activated, $t$ is the top of the curve, and $k$ is the slope factor. Whilst the voltage-dependence of $I_K$ inactivation was fitted with the sum of two Boltzmann equations: $y = (tF / (1 + \exp((_1V_{50} - E) /_1k)) + (t(1 - F) / (1 + \exp((_2V_{50}-E)/_2k)$, where E is the applied voltage, $_1V_{50}$ is the voltage at which 50 % of the 1st component channels are inactivated, $_2V_{50}$ is the voltage at which 50 % of the 2nd component channels are inactivated, $t$ is the top of the curve, $_1k$ is the slope factor for the first component and $_2k$ for the second component, and F defines the relative component contribution.

For current clamp experiments a HEKA EPC-10 amplifier (Lambrecht, Germany) and the corresponding Patchmaster software were used. The extracellular solution contained (in mM): NaCl (140), KCl (4), $MgCl_2$ (1), $CaCl_2$ (2), glucose (4) and HEPES (10), pH 7.40. Patch pipettes, pulled as for $I_K$ experiments, were filled with intracellular solution containing (in mM): KCl (110), NaCl (10), $MgCl_2$ (1) EGTA (1), HEPES (10), $Na_2ATP$ (2), $Na_2GTP$ (0.5), pH 7.3. After gaining access to cells and entering current clamp mode the resting membrane potential of neurons was noted. Ramp depolarisation from 0 pA to 1 nA over a period of 1 s was first used to assess action potential threshold. A step protocol (Δ10 pA, 50 ms) was then used to confirm thresholds. The ability of neurons to fire multiple action potentials was assessed by applying a suprathreshold (2x the threshold determined by step protocol) for 500 ms. Lastly, capsaicin (1 µM in extracellular solution) sensitivity was assessed in voltage clamp mode; neurons that produced an inward current, time-locked to a 5 s application were considered responders. Only cells which fired action potentials and had a resting membrane potential less than or equal to -40 mV were taken through to analyses. Action potential parameters were measured from those evoked by the step protocol using Fitmaster software (HEKA) and IgorPro (Wavemetrics).

### List of Supplemental Figures and Tables

**Study A** DNA Sampling

<u>Inclusion Criteria</u>

Females who are

- Aged 18 years and above
- Able to communicate in English
- Caucasian
- Able to provide written and informed consent
- Who experienced term (beyond 37 week gestation) spontaneous vaginal delivery as nulliparous partituents
- Were healthy during the gestation of the first born

<u>Exclusion Criteria</u>

Females who

- Requested **or** was provided systemic or regional analgesia, including inhalation anaesthetics, spinal or epidurals and opioids (any routes) during delivery of their first child
- Reported having no opportunity for labour analgesia for any reason.
- Required assisted vaginal delivery, including use of Ventouse or forceps for their first child
- Had diabetes or hypertension induced by pregnancy of their first born
- Have known neurological (including channelopathies causing congenital insensitivity to pain) or psychiatric impairments

**Study B** Psychometrics, sensory, pain threshold and tolerance assesments

<u>Inclusion criteria</u>

Females

- who donated DNA in Study A or their corresponding controls

Exclusion criteria

Females who

- are pregnant or breast-feeding
- have any rash, broken skin or skin irregularities where sensory testing is performed
- any underlying medical condition or taking any drug that in the opinion of the investigator will interfere with quantitative sensory testing

**Table S1**. Eligibility criteria for Study A and Study B

| | Test cohort | | | Control cohort | | | | | |
|---|---|---|---|---|---|---|---|---|---|
| **Variable** | n | mean | SD | n | mean | SD | *P* | CI5 | CI95 |
| **Questionnaires** | | | | | | | | | |
| HADS (Anxiety) | 39 | 6.05 | 2.33 | 33 | 6.88 | 3.57 | 0.25845 | -0.62534 | 2.28035 |
| HADS (Depression) | 39 | 2.10 | 1.37 | 33 | 2.48 | 2.18 | 0.77621 | -0.99998 | 0.99996 |
| PCS (Total) | 39 | 9.56 | 6.97 | 33 | 11.18 | 7.50 | 0.41189 | -1.99996 | 5.00002 |
| MHLC (Internal) | 39 | 26.59 | 3.23 | 33 | 26.85 | 3.23 | 0.73576 | -1.26431 | 1.78179 |
| MHLC (Chance) | 39 | 17.31 | 5.40 | 33 | 18.48 | 3.92 | 0.15776 | -0.99993 | 4.00008 |
| MHLC (Powerful Others) | 39 | 14.44 | 4.06 | 33 | 14.97 | 4.65 | 0.60498 | -1.51513 | 2.58273 |
| LOTR (Total) | 39 | 17.46 | 4.60 | 33 | 16.97 | 4.61 | 0.65036 | -2.99994 | 1.99996 |
| **Computerized cognitive assessments (CANTAB)** | | | | | | | | | |
| **Motor Screening Task** | | | | | | | | | |
| Mean latency | 38[#] | 761.3079 | 447.9984 | 30[#] | 687.06 | 135.79 | 0.85586 | -64.69994 | 43.80002 |
| Mean error | 38[#] | 7.208883 | 1.562239 | 30[#] | 7.02 | 1.84 | 0.42846 | -1.16448 | 0.50038 |
| **Rapid Visual Information Processing (RVP)** | | | | | | | | | |
| RVP A' | 36[#] | 0.930904 | 0.056894 | 30[#] | 0.92 | 0.04 | 0.16740 | -0.04706 | 0.00919 |
| RVP B' | 35[#] | 0.890124 | 0.333482 | 29[#] | 0.95 | 0.05 | 0.47025 | -0.01785 | 0.03418 |
| **Spatial Working Memory** | | | | | | | | | |
| Strategy | 38[#] | 27.97 | 8.19 | 30[#] | 30.10 | 6.01 | 0.29844 | -1.00001 | 5.00003 |
| Total errors | 38[#] | 16.39 | 15.99 | 30[#] | 19.67 | 15.51 | 0.22973 | -2.00006 | 10.99998 |
| **Intra-Extra Dimensional Set Shift** | | | | | | | | | |
| Total errors (adjusted) | 37[#] | 18.73 | 16.45 | 30[#] | 19.37 | 16.92 | 0.45651 | -2.00000 | 3.99998 |
| Stages completed | 37[#] | 8.81 | 0.57 | 30[#] | 8.70 | 0.70 | 0.41987 | -0.00003 | 0.00004 |
| Total trials (adjusted) | 37[#] | 83.89 | 29.30 | 30[#] | 84.77 | 29.12 | 0.51143 | -3.99997 | 6.99994 |
| **One Touch Stockings of Cambridge** | | | | | | | | | |
| Mean choices to correct | 37[#] | 1.09 | 0.07 | 30[#] | 1.20 | 0.23 | 0.06084 | -0.00005 | 0.10006 |
| Mean latency to correct | 37[#] | 9689.43 | 4320.31 | 30[#] | 10839.52 | 3998.01 | 0.06460 | -70.24996 | 3115.09998 |

**Table S2** Psychometric results for Study B. HADS, Hospital Anxiety and Depression Scale; PCS, Pain Catastrophising Scale; MHLC, Multi-dimensional Health Locus of Control; Life Orientation Test-Revised (LOTR). n, number of participants; #equipment unavailable/failure; SD, standard deviation; CI5-CI95, 5-95% confidence interval.

| **A** **Pain threshold** | KCNG4+ | | | KCNG4 - | | | P unadjusted | *P* adjusted* | CI5 | CI95 |
|---|---|---|---|---|---|---|---|---|---|---|
| | n | mean | SD | n | mean | SD | | | | |
| Heat (°C) | 3 | 10.1 | 5.00 | 69 | 14.2 | 9.10 | 0.31000 | NA | -17.3 | 9.0 |
| Cold (°C) | 3 | 43.8 | 3.00 | 69 | 43.3 | 3.20 | 0.80000 | NA | -7.2 | 8.2 |
| Cuff-pressure (mmHg) | 3 | 196.2 | 13.80 | 69 | 139.7 | 56.10 | 0.00290 | 0.0090 | 29.7 | 83.4 |

| **B** **Pain threshold** | Test cohort (KCNG4+ individuals excluded) | | | Control cohort | | | P unadjusted | *P* adjusted* | CI5 | CI95 |
|---|---|---|---|---|---|---|---|---|---|---|
| | n | mean | SD | n | mean | SD | | | | |
| Cuff-pressure (mmHg) | 3 | 164.2 | 56.20 | 33 | 113.0 | 9.30 | 0.00008 | 0.0005 | 27.2 | 75.1 |

**Table S3(A)** Effect of the rare allele of *KCNG4* on pain thresholds. *KCNG4+*, individuals who possess the rare allele, *KCNG4-*, controls who do not possess the rare allele; n, number of participants; SD, standard deviation; * Sidak's correction; CI5-CI95, 5-95% confidence interval. **(B)** Comparison of the Test cohort (women who do not possess the rare *KCNG4* allele and did not require analgesic during nulliparous labour) and Control cohort. n, number of participants; SD, standard deviation; * Sidak's correction; CI5-CI95, 5-95% confidence interval.

| A | $K_V$6.4 | $K_V$6.4-Met419 |
|---|---|---|
| *n* | 8 | 7 |
| Capacitance (pF) | 22.9 ± 1.4 | 23.2 ± 1.8 |
| Access resistance (MΩ) | 8.2 ± 1.1 | 7.8 ± 0.9 |
| *Activation* | | |
| $V_{1/2}$ (mV) | -5.4 ± 1.8 | -9.8 ± 1.1 |
| *k* | 8.6 ± 1.5 | 8.9 ± 0.9 |
| *Inactivation* | | |
| 1st component | | |
| $V_{1/2}$ (mV) | -0.8 ± 29.5 | -36.2 ± 3.3 |
| *k* | -46.1 ± 25.6 | -63.9 ± 26.5 |
| 2nd component | | |
| $V_{1/2}$ (mV) | -60.2 ± 6.6 | -33.8 ± 2.1** |
| *k* | -29.6 ± 8.1 | -26.4 ± 15.6 |

| B | $K_V$6.4-Met419 | $K_V$6.4 | Untransfected |
|---|---|---|---|
| n | 10 | 8 | 6 |
| RMP (mV) | -50.10 ± 2.05 | -47.33 ± 1.14 | -46.00 ± 2.14 |
| Capacitance (pF) | 41.54 ± 10.17 | 31.53 ± 3.69 | 21.55 ± 4.78 |
| Ramp Threshold (pA) | 248.60 ± 50.33* | 91.56 ± 16.74 | 112.50 ± 32.51 |
| Number of ramp AP | 12.70 ± 3.48 | 10.22 ± 2.47 | 15.50 ± 4.79 |
| Step Threshold (pA) | 172.00 ± 34.44* | 61.11 ± 12.18 | 88.33 ± 13.76 |
| Amplitude (mV) | 75.20 ± 5.20 | 76.66 ± 6.72 | 59.87 ± 4.62 |
| HPD (ms) | 3.79 ± 0.81 | 5.69 ± 1.26 | 3.71 ± 0.62 |
| AHP Duration (ms) | 16.48 ± 1.65 | 31.53 ± 7.35 | 17.17 ± 3.16 |
| $AHP_{50}$ (ms) | 8.52 ± 0.75 | 10.32 ± 2.15 | 8.75 ± 1.40 |
| AHP Amplitude (mV) | 18.49 ± 1.80 | 15.92 ± 1.97 | 17.95 ± 2.90 |
| AP Freq @ 2xThr | 6.40 ± 1.17 | 3.00 ± 0.73 | 4.33 ± 2.44 |

**Table S4 (A)** Electrophysiological characteristics of mouse sensory neurons transfected with wild-type $K_V$6.4 and $K_V$6.4-Met419. ** $P < 0.01$ **(B)** Action potential parameters of mouse sensory neurones transfected with wild-type KV6.4 or $K_V$6.4-Met419 and untransfected cells from current clamp experiments. RMP, resting membrane potential, AP, action potential, HPD, half peak duration, AHP, afterhyperpolarisation duration, Thr,, threshold, Freq., frequency. *P <0.05

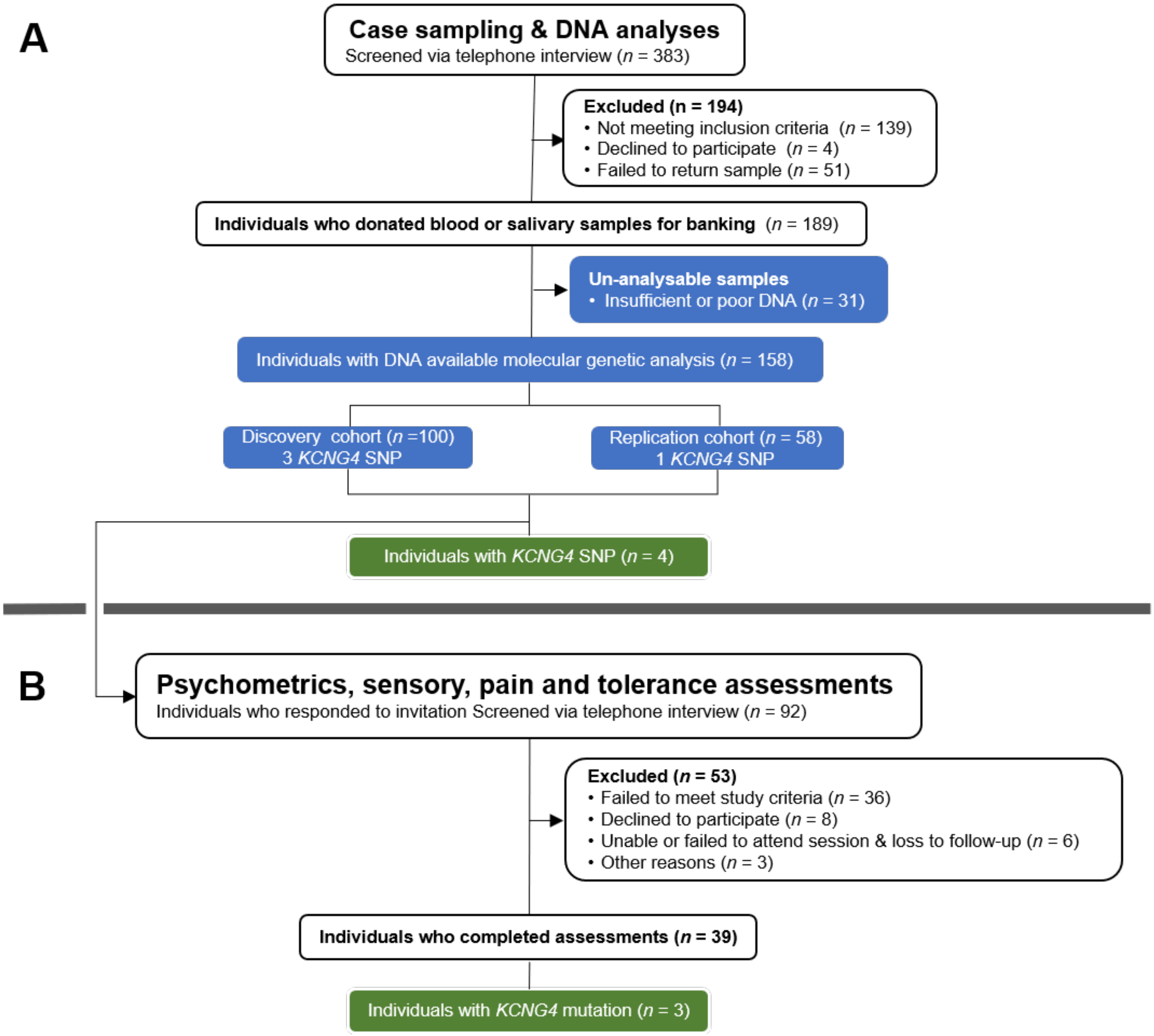

**Figure S1** Flow-chart illustrating the recruitment and screening of participants for (A) genetic sampling and (B) the subset of those participants who underwent psychometric, sensory and pain (threshold and tolerance) assessments. Blue rectangles indicate handling, processing and analyses of DNA samples that were donated by participants. Green rectangles indicated number of individuals assessed or DNA analysed with *KCNG4* mutation. n, number of samples or individuals.

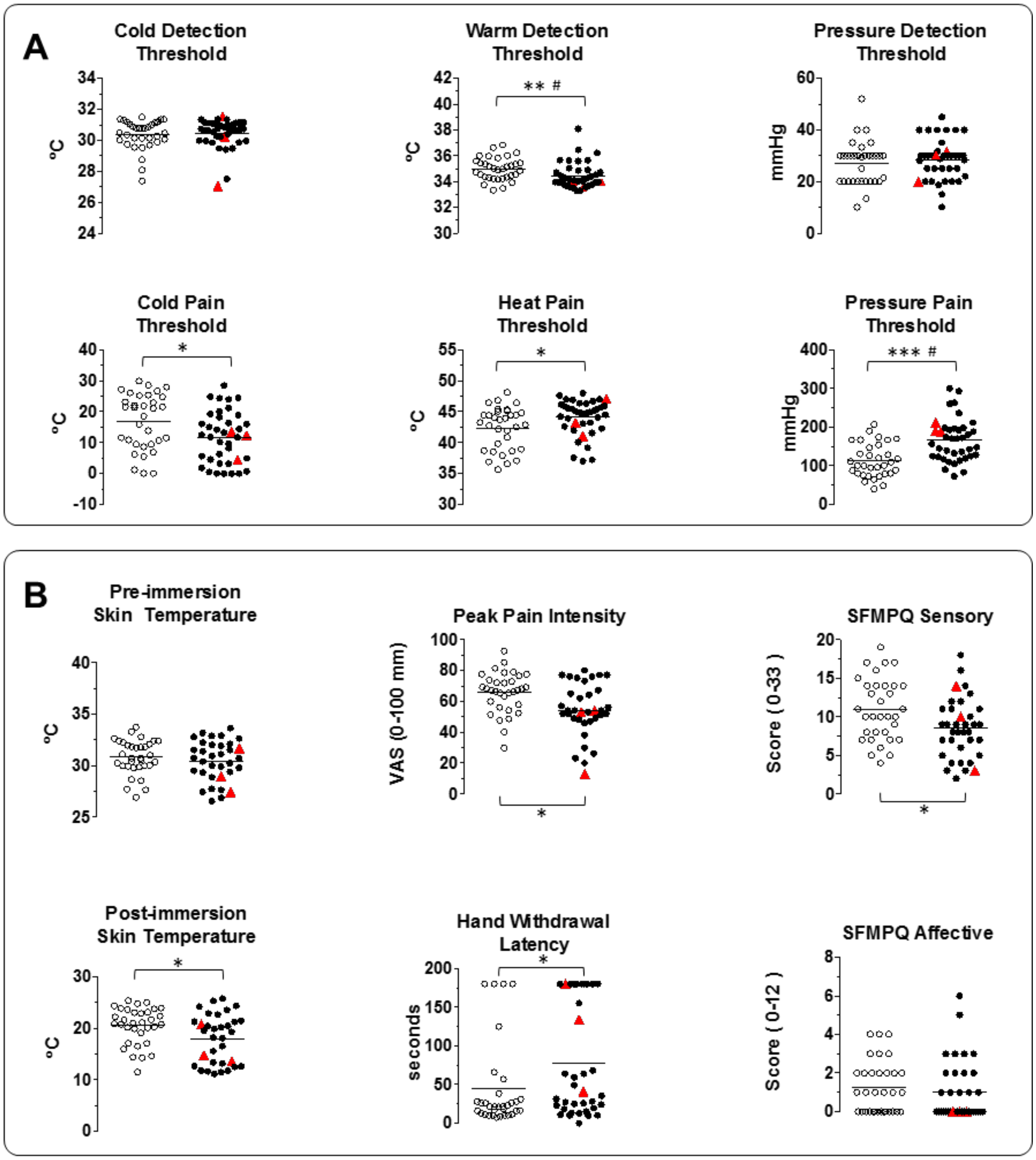

**Figure S2** Sensory detection, pain threshold and tolerance assessments (A) Thresholds for sensory detection and pain for heat, cold and cuff pressure. (B) Testing of pain tolerance to hand immersion in cold water. Left-sided graphs: skin temperatures pre- and post-immersion. Middle graphs: withdrawal latency and ratings of peak pain experienced during hand immersion. Bottom graphs: ratings of the sensory and affective qualities of pain experienced with the SFMPQ. Clear circles indicate individuals in control cohort, and filled circles indicate those in the test cohort. The three individuals with KV6.4 p.Val419Met are indicated by red triangles. Horizontal lines represent the mean for each cohort. * P<0.05, ** P<0.01 *** P<0.001; # Sidak adjusted P <0.05

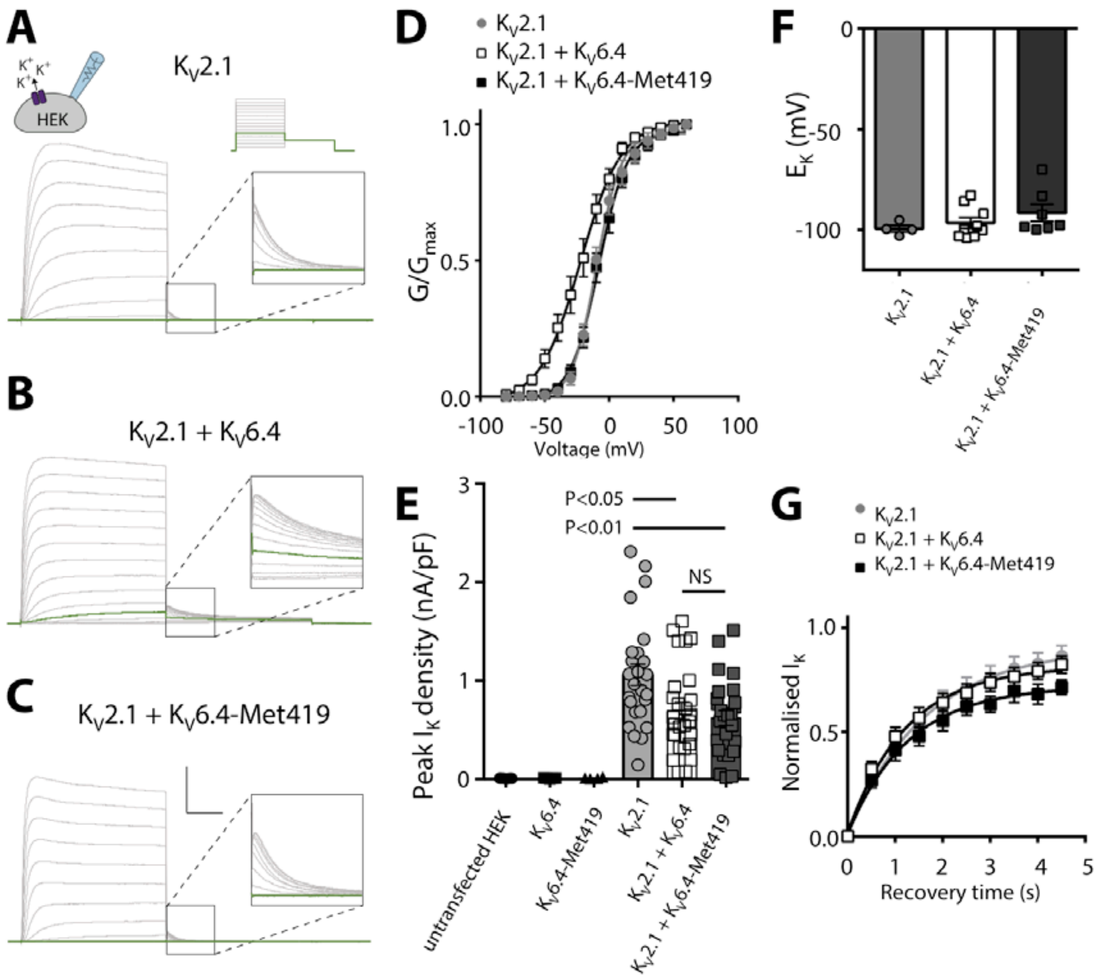

**Figure S3** Supporting electrophysiology data for the functional effects of $K_V6.4$ and $K_V6.4$-Met419 on $K_V2.1$ currents in HEK293 cells Representative current recordings to determine $K_V2.1$ (A), $K_V2.1/K_V6.4$ (B), $K_V2.1/K_V6.4$-Met419 (C) channel activation properties. The applied voltage protocols are illustrated above the currents shown in (A). Vertical scale bar is 10 nA, horizontal scale bar is 50 ms. Green traces indicate currents recorded during the −40 mV prepulse. D. Voltage-dependence of activation of $K_V2.1$ (grey filled circles, $n$ = 13), $K_V2.1/K_V6.4$ (open squares, $n$ = 14), and $K_V2.1/K_V6.4$-Met419 (black squares, $n$ = 13). The voltage-dependence of activation was determined by normalising tail currents at −60 mV as a function of a prepulse from −80 to +60 mV, in +10 mV increments. Solid lines represent the Boltzmann fitted curves. (E) Peak $K^+$ current density obtained from +30 mV step of voltage protocol. Bars indicate mean values, error bars indicate SEM. First three groups, $n$ = 4-7 from 2 independent experiments, last three groups, $n$ = 25-27 from 5 independent experiments. (F) Reversal potential obtained from a linear fit of tail currents from −10 mV to a series of voltage steps from −140 to −50 mV, in +10 mV increments. Bars indicate mean values, error bars indicate SEM, $n$ = 4-9. (G) Recovery time from inactivation of $K_V2.1$ (grey filled circles, $n$ = 6), $K_V2.1/K_V6.4$ (open squares, $n$ = 10), and $K_V2.1/K_V6.4$-Met419 (black squares, $n$ = 9). Relative peak current plotted from a 200 ms test pulse to +20 mV at various time intervals following a 5 s prepulse to +20 mV. Solid lines represent exponential fitted curves.

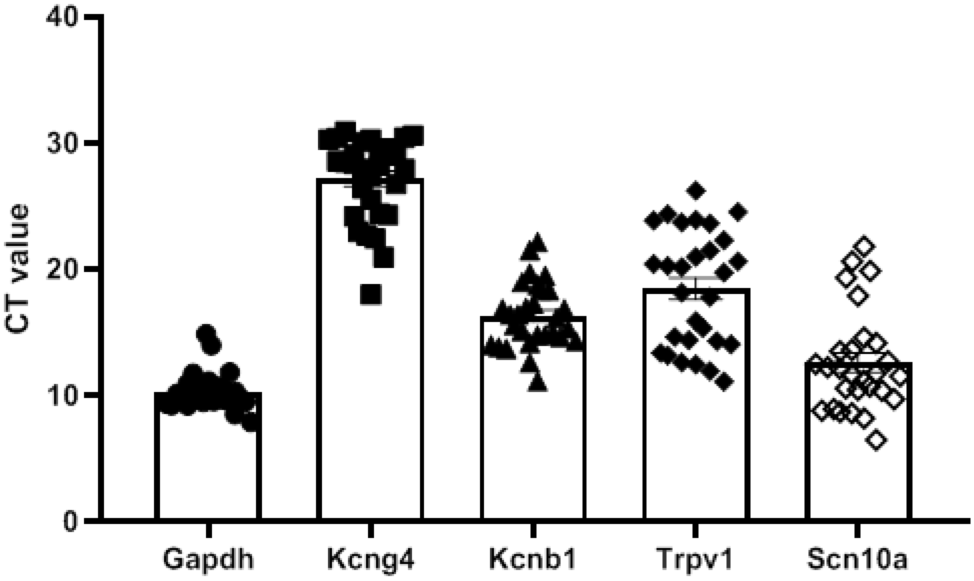

**Figure S4** Graph demonstrating the mean raw cycle threshold (CT) values of *Kcng4*-positive mouse uterus-innervating sensory neurones for each gene assessed by single cell quantitative PCR analysis. Error bars represent standard error mean.

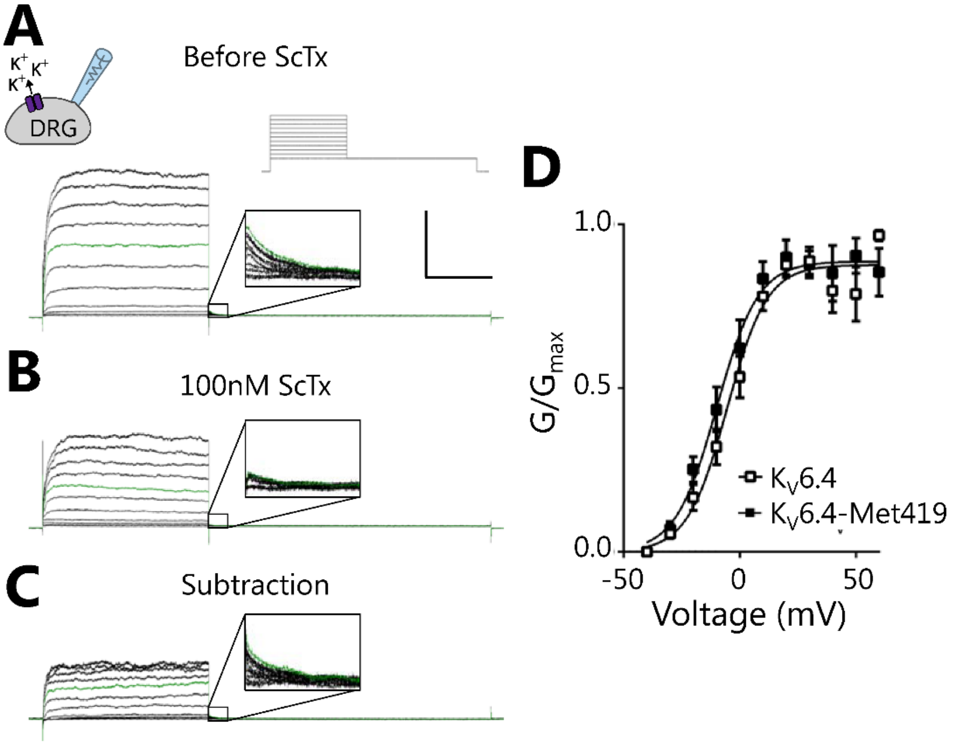

**Figure S5** Effect of $K_V$6.4 and $K_V$6.4-Met419 on the voltage dependence of activation of the stromatoxin-1-sensitive current in mouse sensory neurons (A) Representative $I_K$ recordings produced by *inset* voltage protocol in the absence and presence of 100nM ScTx (B). (C) The ScTx-sensitive $I_K$ is isolated by subtraction of B from A. Expanded tail currents are shown for all three representative traces, each *inset* is 50 ms by 450 pA. The green tracing in A, B and C represent the current at +20 mV. (D) Activation curve of the ScTx-sensitive $I_K$ obtained from mouse sensory neurons transfected with either wild-type $K_V$6.4 or $K_V$6.4-Met419. In both cases a Boltzmann function was fit to the data.

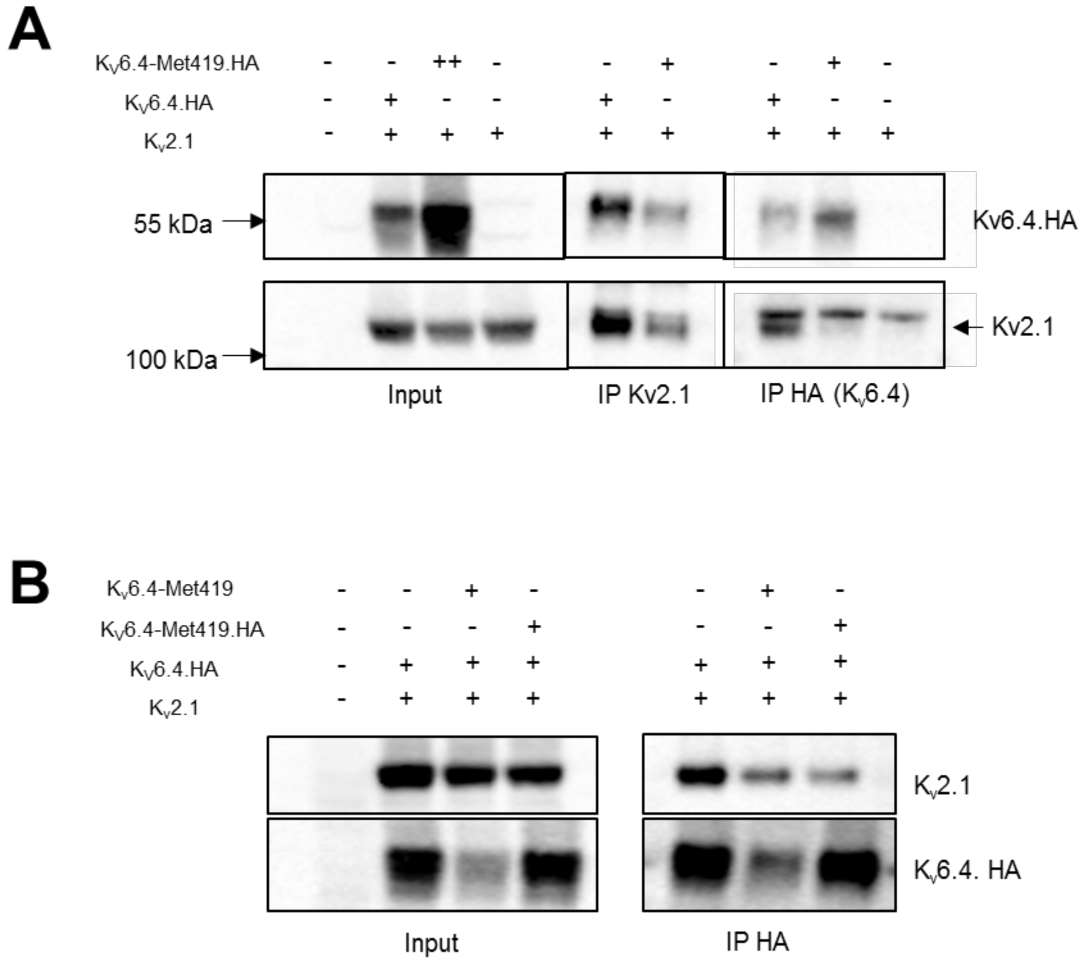

**Figure S6 (**A) Co-IP experiments showing absence of $K_V2.1$ binding to $K_V6.4$-Met419 is not due to $K_V6.4$-Met419 lack of stability. There is significantly reduced binding of $K_V6.4$-Met419 to $K_V2.1$ even when significantly overexpressed compared to $K_V6.4$. This blot also confirms that HA antibody does not pull down $K_V2.1$ in the absence of $K_V6.4$ expression. (B) Co-immunoprecipitation experiment for $K_V6.4$ and $K_V2.1$ demonstrating that there is similar reduced binding for $K_V6.4$ to $K_V2.1$ in the heterozygous mutant state, whether or not the $K_V6.4$-Met419 is tagged with HA or not.

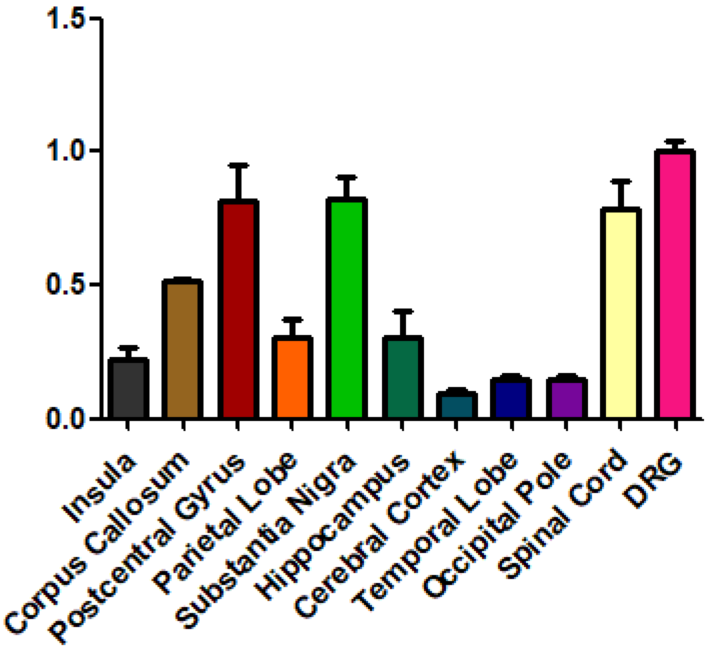

**Figure S7** Expression levels of *KCNG4* in different human brain regions, the spinal cord and DRG (dorsal root ganglion). The graph displays the mean of three mRNA/cDNA conversions, assessed by TaqMan qPCR normalised to a *GAPDH* control and compared with the highest expressing tissue, the DRG.
